# Curated annotations locate biological concepts in protein language models from layers to neurons

**DOI:** 10.64898/2026.11.19.767884

**Authors:** Logan Hallee, Jason P. Gleghorn

**Affiliations:** Center for Bioinformatics and Computational Biology, University of Delaware; Department of Biomedical Engineering, University of Delaware; Synthyra, PBLLC

**Keywords:** Interpretability, Protein language models, Probing, Neurons, Sparse autoencoders, Protein-protein interaction

## Abstract

Interpretability methods describe what a network has learned, yet their claims are rarely tested against model-independent ground truth. We argue that proteins are an ideal modality for interpretability since curators annotate functional sites residue by residue. Across three ESMC protein language models (300M-6B parameters), we show that conclusions depend on where models are read, how concepts are decoded, which units are examined, and how they are scored. Concepts become linearly readable along depth long before erasure impairs masked-token prediction, and 20-fold scaling shifts concept depth more than peak score. Most of a nonlinear probe’s lead over a linear one reflects fitting and readout shape rather than nonlinear encoding. Over whole proteins, saturated metrics render feed-forward neurons indistinguishable from random rotations. At annotated residues, dual-signed neuron poles identify 2.5-5.4 times as many concepts as rotations without training, though sparse autoencoders identify more. Without held-out validation, however, shuffled annotations label nearly every latent, and causal reliance emerges in neuron groups rather than single poles. Reading selected intra-block states exposes binding and active sites that stream probes read poorly and lifts a simple probe 5% above the best published MCC on a leakage-controlled interaction benchmark. We recommend reading a model directly against annotation, scoring units where they fire with a measure that is not saturated, and training a dictionary only where it adds coverage.

## Introduction

Protein language models (pLMs), typically transformer neural networks trained by masked language modeling (MLM) on protein sequences, learn biological associations by compressing sequence data into their weights(1–3). Their representations drive protein property prediction and design tasks across life science domains(4–11). Unfortunately, we understand how pLMs organize information only in part. Various probing studies have traced the depth at which layer outputs correlate with a concept(12–17), while sparse autoencoders (SAEs), dictionaries of sparsely active latents trained to reconstruct a hidden state, have been used to extract precise features(18–21). InterPLM associated up to 2,548 SAE latents per layer of ESM2-8M with Swiss-Prot concepts, against at most 46 neurons(22). Candido and colleagues have since trained dictionaries on every ESMC block, and one is released with descriptions of each of its 16,384 latents written by a multi-LLM-agent system(23, 24).

It is less clear which of a model’s dense states and units hold a concept. Each block computes queries, keys, values, and feed-forward activations, yet probes have read feed-forward neurons or attention heads one type at a time(25–29). The neurons that InterPLM compared with SAE latents were dimensions of the residual stream, where no elementwise nonlinearity acts(22). An SAE is itself a trained model, one for each layer that it reads, and its latents are usually named by a language model(24, 30). These names are rarely checked against an independent record of what a latent should respond to, although random baselines nearly match trained SAEs on standard interpretability scores in language models(31). The score is part of the problem, since a unit scored by whether it fires in the proteins that carry its term sits on a scale where almost any direction succeeds.

Recent work in vision and language argues that dense SwiGLU feed-forward neurons are largely interpretable once each is read as two poles, one for each sign of its output, and that communities of neurons spread across layers carry behaviors that no single neuron does(32–34). We argue that pLMs are an ideal testing ground for such claims. Proteins are a valuable language modality, and they allow a more direct reading than natural language because Swiss-Prot curators annotate active sites, binding sites, motifs, and domains residue by residue (token by token), leaving an opportunity for precise attribution. Controlled concept datasets for supervised probes can be processed with sequence homology-aware splits to prevent obvious data leakage(14, 35).

We read ESMC models against curated annotation from global to local(6, 23). Globally, linear and multilayer perceptron (MLP) probes read every token-wise state of each model (453 to 1,203 states) on the 22 concept datasets of Whitfield and colleagues(14). Between the two scales, we discovered communities of feed-forward neurons that act together across blocks. Locally, we scored each feed-forward neuron, read as two poles, on 636 Swiss-Prot concepts annotated by residue, compared to random rotations, residual stream dimensions, and SAE latents.

Here we show that a conclusion at each of these scales depends on where a concept is read, how it is decoded, which units are read, and how they are scored. Concepts become linearly readable in an order set by their kind, long before the model depends on them, and most of a nonlinear probe’s lead comes from fitting. States inside a block expose binding and active sites that linear stream probes read poorly, and communities of neurons carry predictions that no single neuron carries. Single neurons mark curated concepts far more often than random directions, which only a residue-level score reveals, although SAEs mark more. The state these checks select lets a simple probe of a frozen ESMC-6B exceed published sequence-based predictors on a gold-standard interaction benchmark.

## Results

We studied ESMC-300, ESMC-600, and ESMC-6B, with 30, 36, and 80 blocks, respectively (**Fig. 1**). The token embedding, each block’s output, and the final layer norm form the residual stream, which we map to a relative depth axis from 0 at the token embedding to 1 at the final layer norm(36). Our analysis read 18 types of latent states, including 14 computed inside a block and ESMFold2’s structure stream, yielding 453, 543, and 1,203 states in the three models (**Supplementary Table 3**). The 22 concept tasks span composition, residue sites and modifications, sorting and topology, InterPro membership, function, and secondary structure, and 13 of them label whole proteins(14, 37). Probe scores are the MCC unless stated otherwise, and *x ± h* gives half the width of a 95% interval over the test proteins(38).

**Fig. 1.**
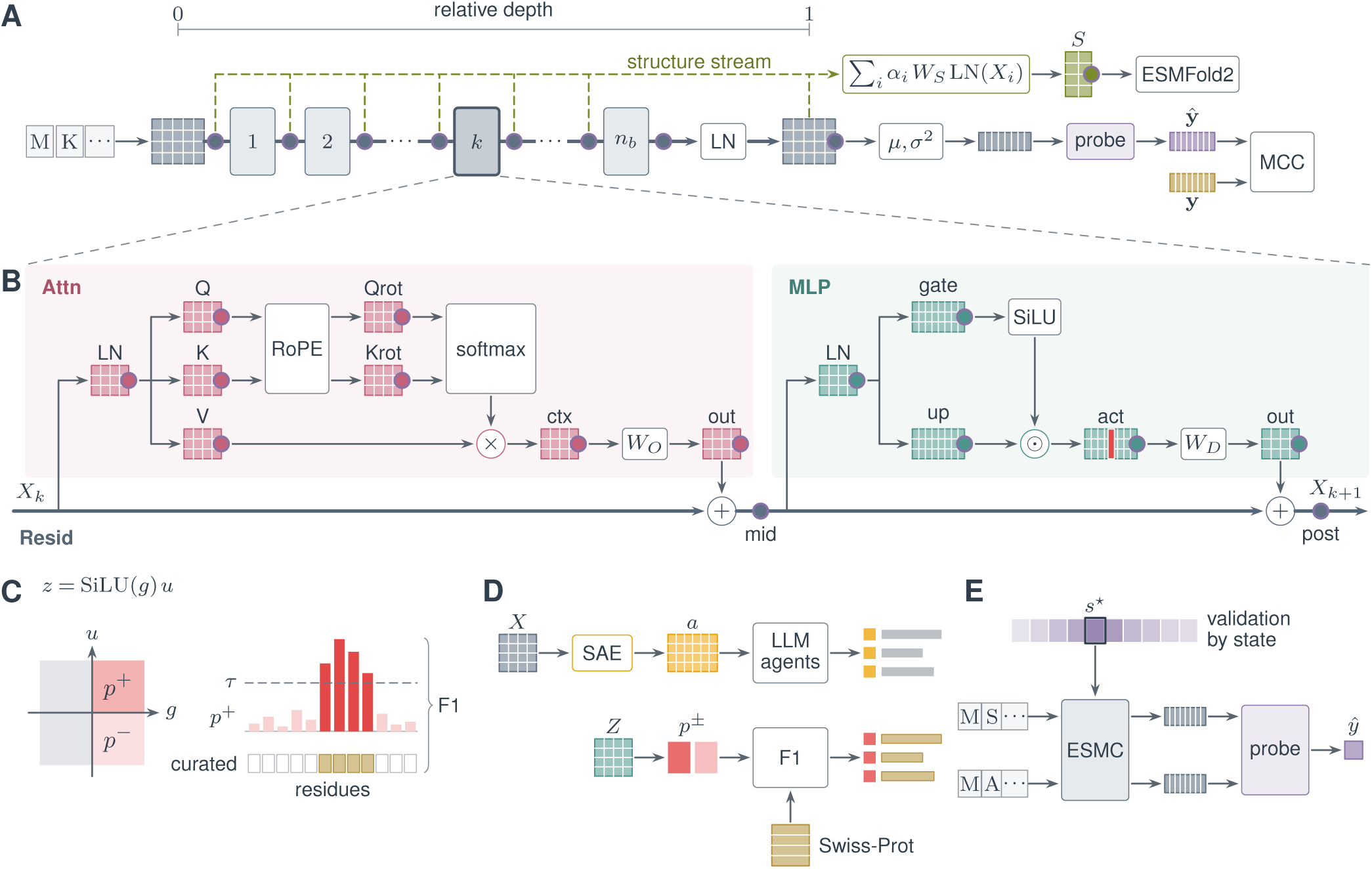
Reading ESMC from global to local, and using what is read. **A**, a protein’s tokens pass through *n_b_* blocks and a final layer norm (LN), and every ringed state of the residual stream is probed, from relative depth 0 at the token embedding to 1 at the final norm. A probe, drawn out at the final norm, pools a state over the protein’s residues by its mean *µ* and variance *σ*^2^, reads it with a linear or one-hidden-layer MLP readout, and scores it by MCC against curated labels **y**. The structure stream *S* that ESMFold2 reads is probed as one more state; it is a weighted sum of the stream’s states, each passed through one shared layer norm and projection *W_S_*. **B**, block *k* of **A**. We probe every token-wise state, covering the states of attention, the stream between the branches and after the block, and the states of the SwiGLU feed-forward network, in all 453, 543, and 1,203 states in ESMC-300, ESMC-600, and ESMC-6B once the structure stream of **A** is counted. **C**, one feed-forward neuron, *z* = SiLU(*g*) *u*, read as two poles, *p*^+^ and *p^−^*, the positive and negative output with the gate open (*g >* 0), outputs of a closed gate being left out. A pole is active along a protein’s residues where its activation exceeds a threshold *τ*, and these residues are scored by F1 against those that curators annotated with a concept. **D**, two routes from a dense model to named units. In the first, a sparse autoencoder (SAE) trained on a layer’s stream *X* gives latents *a*, which a multi-agent language-model system describes in free text. In the second, the poles *p^±^* of the feed-forward product *Z* are scored by F1 against Swiss-Prot’s curated annotation vocabulary and take its terms, with no training and no language model description. **E**, the state *★* that reads a task best on validation is read from ESMC for two proteins, and their pooled states feed one probe that predicts whether they interact. Matrices, bars, and cells are schematic, not measured values.

### A concept is readable long before the model uses it

We first asked where along pLM depth each kind of concept becomes readable. A linear probe read every stream state for each task, and a task’s onset is the first depth at which its score comes within 5% of its highest (**Fig. 2A**). Composition needs no depth, as the probe on each model’s token embedding already reaches 96% of its highest score. Secondary structure comes last in every model, with onsets at a relative depth of 0.81 or later. Between these ends, the order shifts with scale: InterPro memberships move earlier, from 0.42 in ESMC-300 to 0.30 in ESMC-6B, while the other three kinds move later.

**Fig. 2.**
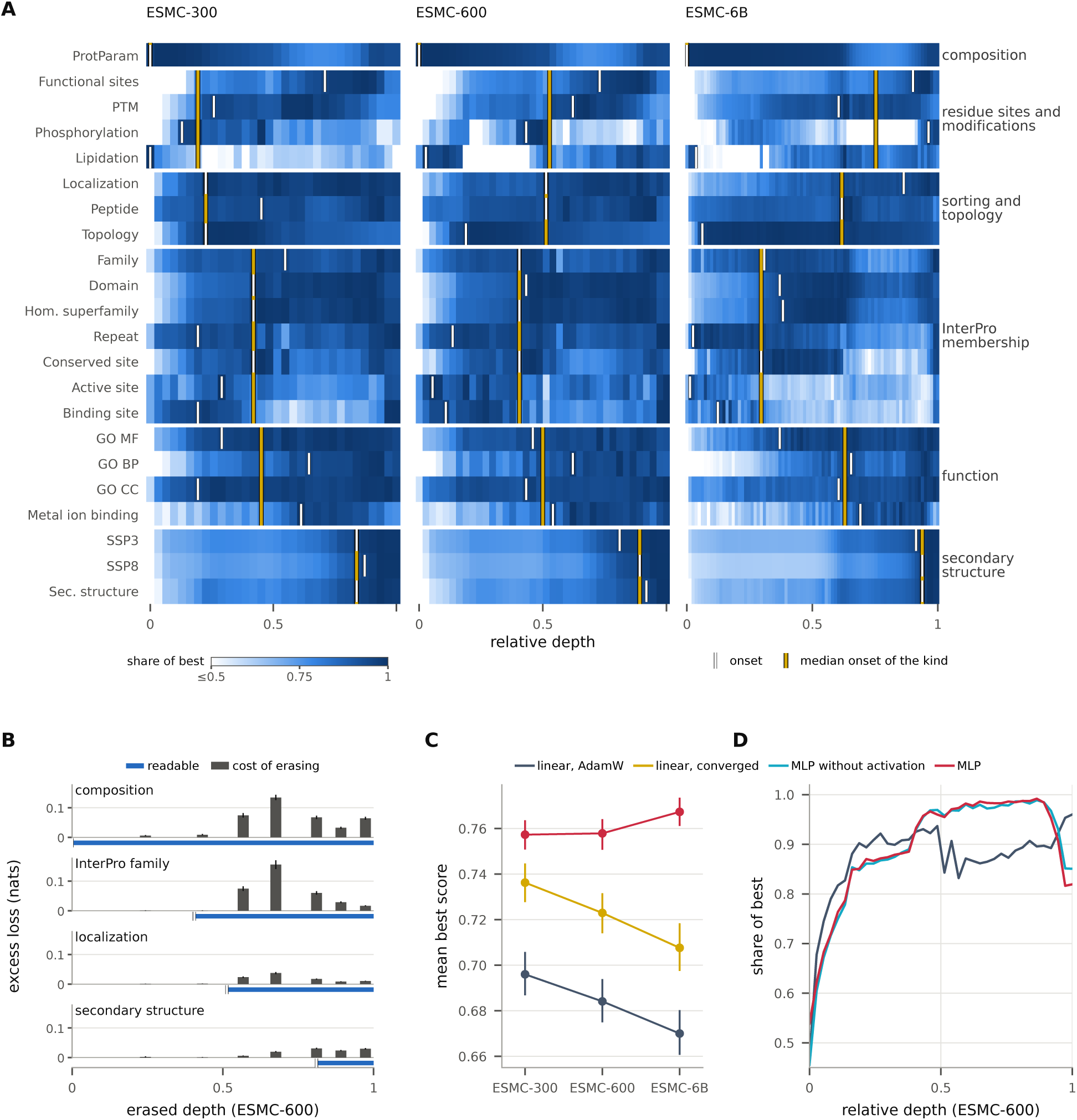
Concepts become readable by kind along depth, long before the model depends on them, and fitting explains much of a nonlinear probe’s lead. Relative depth runs from 0, the token embedding, to 1, the final layer norm. **A**, each task’s residual-stream score as a share of its highest, for the 22 tasks grouped by kind, with each task’s onset (the first depth within 5% of its highest score) and each kind’s median onset marked. **B**, for four concepts in ESMC-600, the masked-token loss that erasing the concept at one depth costs beyond erasing as many random directions, above a band that starts where the concept becomes linearly readable. The first erased block is not drawn, as random directions there cost more than the erasure (**Supplementary Information, page 21**). **C**, mean best score over the 13 protein tasks under AdamW linear probes, converged linear fits, and MLP probes. **D**, InterPro membership along ESMC-600’s stream, as a share of each task’s best score averaged over seven tasks, under the linear probe, the MLP without its activation, and the MLP. Whiskers in B and error bars in C are 95% intervals over 1,000 bootstrap draws of the test proteins.

The 20-fold increase in pLM parameters changes where a concept is read far more than how well. Over the 22 tasks, the mean best stream score is 0.634, 0.629, and 0.626 in ESMC-300, ESMC-600, and ESMC-6B. Yet ESMC-6B reads two kinds far deeper: sorting and topology at a median relative depth of 0.65, and residue sites and modifications at 0.95, versus 0.29 and 0.55 in ESMC-300 (**Supplementary Information, page 18**).

A concept is also readable well before the model depends on it. We tested this in ESMC-600 with LEACE, erasing from every token at one depth what a linear map reads of a concept, against a control that erases as many random directions(39). InterPro memberships are readable from a median relative depth of 0.41, yet erasing family, composition, or localization before 0.5 raises the model’s loss on masked tokens by under 1% more than the control does. At 0.68, the same erasure raises that loss by 11% for family. A linear probe reads the erased concept again by the final norm (**Fig. 2B** and **Supplementary Information, page 21**). A concept’s onset is therefore neither where the model depends on it nor evidence that it is computed there.

### Fit a linear probe to convergence and the nonlinear advantage mostly disappears

We next asked whether a linear probe is the right instrument for this map, because a nonlinear probe’s lead over a linear one often signals that a concept is present but not linearly encoded. Over the 13 protein tasks, MLP probes exceed linear probes trained by AdamW with early stopping, a usual practice, by 0.061 *±* 0.008, 0.074, and 0.097 in ESMC-300, ESMC-600, and ESMC-6B(40, 41). Against linear fits refitted to convergence, the same MLP probes lead by only 0.021 *±* 0.007, 0.035 *±* 0.008, and 0.060 *±* 0.009 (**Fig. 2C**). A randomly initialized ESMC-300 shows no surviving lead (*−*0.008 *±* 0.010), so this occurs from what pretraining put into the weights.

Removing the MLP’s activation leaves a low-rank linear map through a hidden layer that a task’s labels share, trained like the MLP. In ESMC-600, on the pooled mean alone, this map recovers 84% of the MLP’s lead over the AdamW probe, so the activation itself adds only 0.012 *±* 0.004 (**Supplementary Information, page 22**). Along the stream, the linear map follows the MLP through the middle depths, where it stays at 96% or more of its best InterPro score while the per-label linear probe falls to 83% (**Fig. 2D**). A linear probe’s failure at some depth is therefore weak evidence that a concept is absent there. The readout also changes what scale appears to do. Converged linear fits read less from ESMC-6B than from ESMC-300 (0.708 against 0.736), while MLP probes read slightly more (0.767 against 0.757). The input’s treatment can also decide an intervention test, since in a phosphomimetic scan standardized inputs let even a randomly initialized ESMC-300 model separate mutants that keep a phosphosite’s function from those that remove it(14) (**Supplementary Information, page 23**).

### States inside a block expose sites that linear stream probes read poorly

Between depth and single neurons lies the type of state, and we asked whether the states a block computes before it writes to the stream contain anything the stream does not. Compared with the best stream state for each kind of concept, most of the 14 state types inside a block read no better, and they fall into groups that recur at every model size (**Fig. 3A**).

**Fig. 3.**
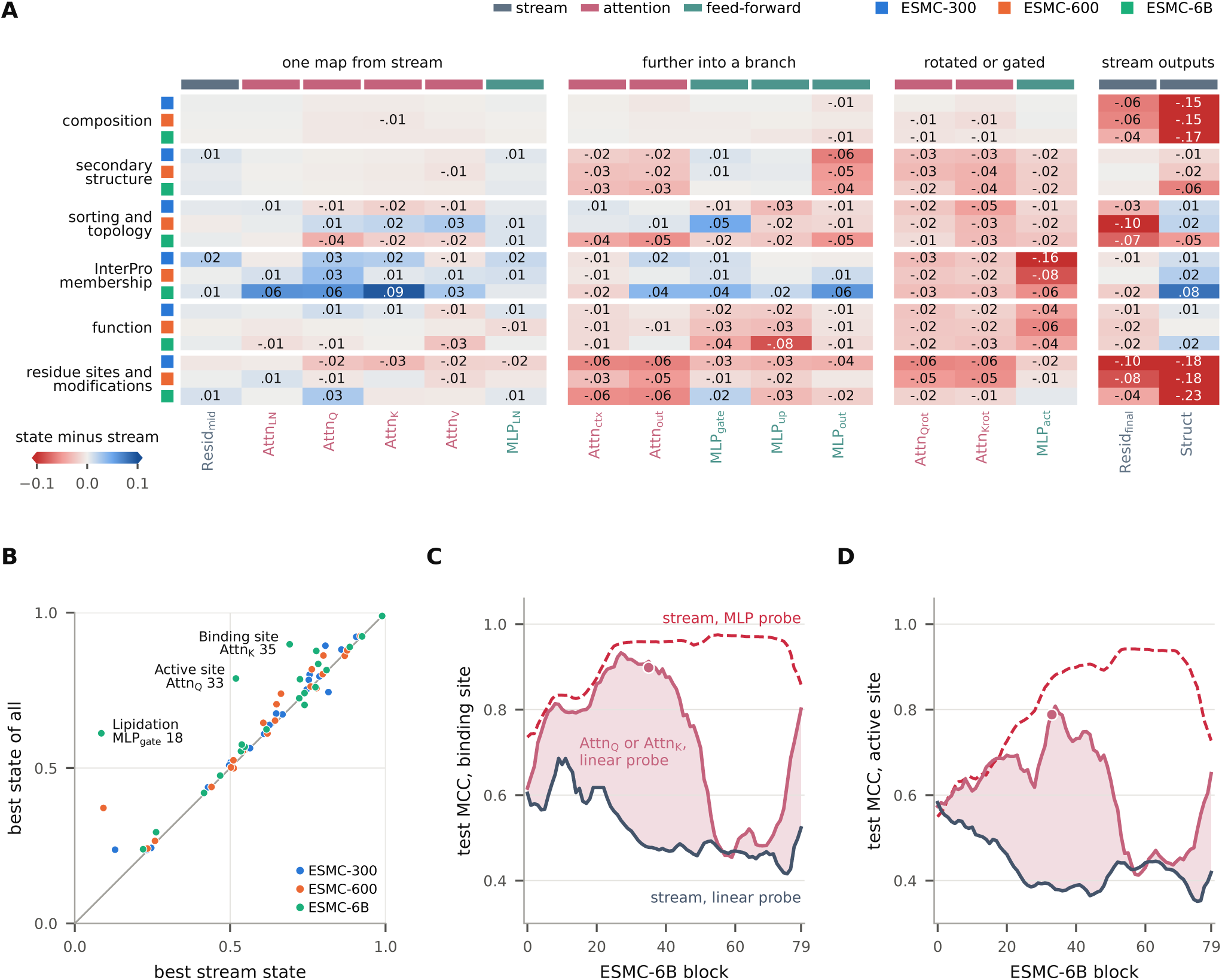
Most states inside a block track the residual stream, and the queries and keys of attention in ESMC-6B expose sites that a linear probe of the pooled stream reads poorly. **A**, each state type’s best probe over blocks minus the task’s best residual-stream probe, as the median over a kind’s tasks, for each kind of concept (rows, one per model) and state type (columns, grouped by how far a state lies from the stream). Values that round to zero are blank. **B**, each task’s best state of all against its best stream state, both chosen on validation, for the 22 concepts. The three tasks of ESMC-6B whose best state leads by 0.1 or more are named with that state. **C**, binding sites and **D**, active sites along ESMC-6B’s blocks, read by an MLP probe of the stream, by a linear probe of the queries or keys of attention, whichever scores higher on validation at each block, and by a linear probe of the stream. The shaded gap is where attention’s projections lead the stream under a linear probe, and the dot is the state named in B. Lines are running means over five blocks.

The best state of a task nonetheless usually lies inside the block, by a median of at most 0.013, and the gains concentrate in ESMC-6B and a few tasks (**Fig. 3B**). There, the state chosen on validation reads binding sites at 0.90 (Attn_K_ of block 35) and active sites at 0.79 (Attn_Q_ of block 33), at least 0.41 and 0.39 above the stream entering or leaving that block. The stream holds these sites equally, as MLP probes on the same blocks read binding sites at 0.96 and active sites at 0.86 to 0.89. The attention projections therefore expose linearly what a linear probe of the pooled stream reads poorly (**Fig. 3C,D**). Most of this advantage comes from probe fitting rather than the state’s content, since refitting to convergence lets the best stream state match or beat the best state inside the block on 7 of the 13 protein tasks (**Supplementary Table 2**). Only the largest model keeps an advantage under converged and tuned fitting, as ESMC-6B’s chosen state inside the block leads its best stream state by 0.035 to 0.049 ROC AUC on interaction under every probe we fitted, against at most 0.011 in the smaller models (**Supplementary Information, page 29**).

### Communities of neurons carry predictions no neuron carries alone

Erasure separated a concept’s readability from the model’s dependence on it, so we next asked which units a masked-token prediction depends on. Between a block’s states and its single neurons, groups of neurons can act together across blocks. We scored how much each feed-forward neuron of ESMC-600 contributes to predicting a masked token, linked the neurons that contribute together, and clustered the links into 39,260 communities of 2 to 128 neurons (**Fig. 4A**). A community explains a held-out prediction when removing its neurons changes the prediction and keeping them alone preserves it. A discovered community explains 2.6% of 38,985 held-out predictions, about 260 times the share that shuffled communities of the same size in each block explain (**Fig. 4B**).

**Fig. 4.**
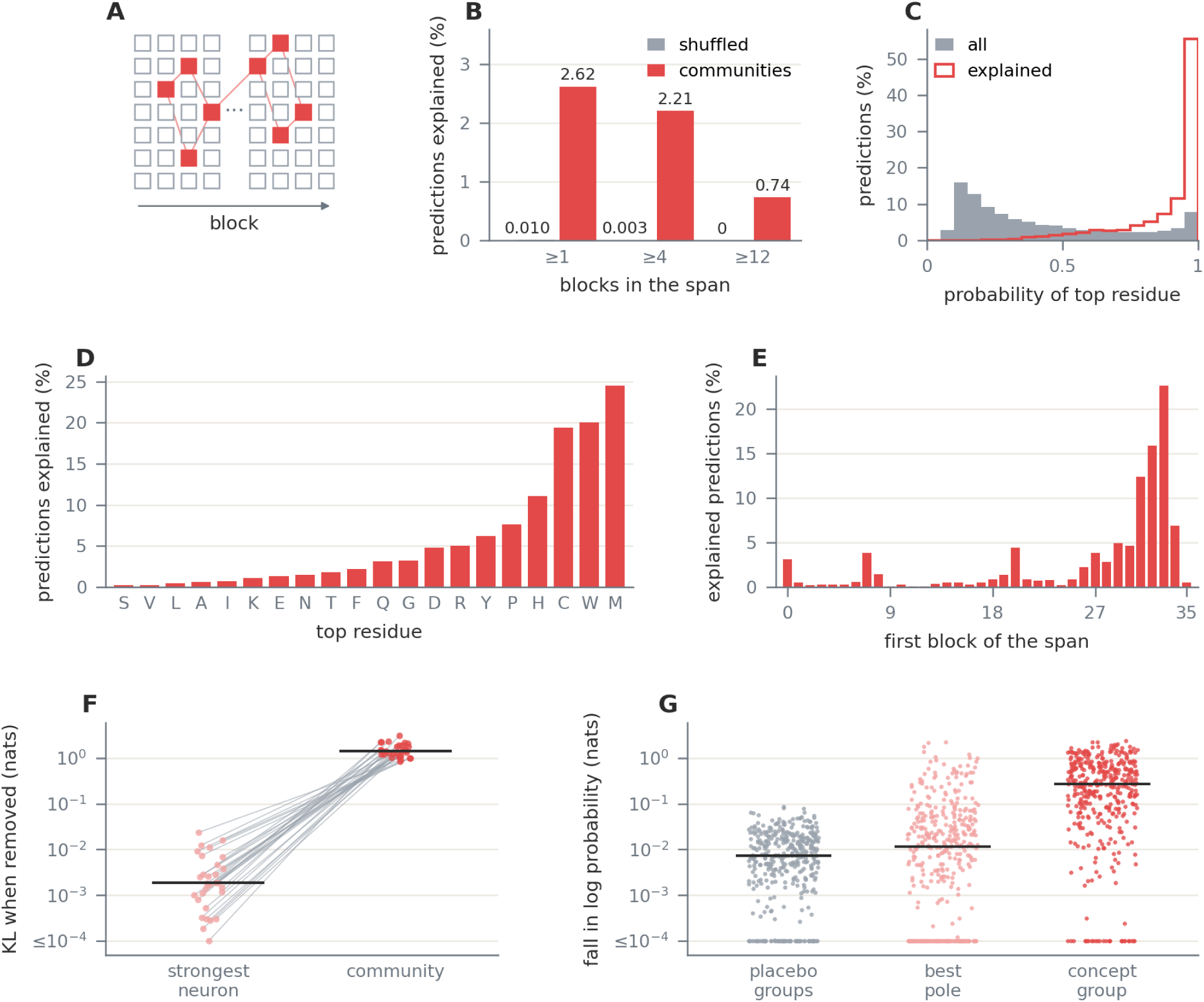
Communities of feed-forward neurons explain the predictions that ESMC-600 is most sure of, and no neuron carries one alone. **A**, a community is a set of neurons in several blocks that contribute together to the model’s predictions. **B**, the share of 38,985 held-out predictions of a masked token explained within a span of at least 1, 4, and 12 blocks by shuffled communities, which keep each community’s size in each block and redraw its neurons, and by discovered communities. **C**, the probability that the intact model gives its top residue, at all predictions and at the 1,022 that a community explains (medians 0.31 and 0.96). **D**, the share of predictions explained by the model’s top residue. **E**, the predictions explained by the first block of the narrowest span of blocks that explains each, which is block 27 or later for 74% of them. **F**, for each of the 30 communities that explain the most predictions, the change in prediction (KL divergence) when its most influential neuron is removed alone (median 0.0018 nats) versus when the whole community is removed (median 1.42 nats), with a line joining each community’s two values. **G**, for 406 Swiss-Prot concepts, the fall in the log probability of the true residue at the concept’s masked tokens when placebo groups of matched activation are removed (median 0.0074 nats, each the mean of ten), when its best single pole is removed (0.0116 nats), and when the poles labeled with the concept across blocks are removed together (0.270 nats). These concept groups are assembled from curated labels, whereas the communities of A to F are found from the model’s predictions. Ticks in F and G are medians, and values at or under 10*^−^*^4^ are drawn at 10*^−^*^4^.

The predictions explained are those the model is most sure of, with a median top probability of 0.96 against 0.31 over all predictions, and they include a quarter of the predictions of methionine and a fifth of those of tryptophan and cysteine (**Fig. 4C,D**). The neurons that explain a prediction sit mostly in the last blocks, and none matters alone, since removing a community moves the prediction about 800 times as far as removing its most influential neuron (**Fig. 4E,F**). In each of the 30 communities that explain the most predictions, no curated term annotates even 20% of the explained residues, so they predict amino-acid identity, not annotated concepts.

### Only a residue-level score separates neurons from random directions

At the local scale, we asked whether a single neuron marks a biological feature at the residues where curators annotated it. In ESMC the MLP’s leverage SwiGLU activation, so each neuron multiplies a gate *g* and an up projection *u* computed from a residue’s state, as *z* = SiLU(*g*) *u*, so one neuron can fire with either sign(42). Following Cammarata, we read it as two units, a positive and a negative pole, each active only when the gate is open(32) (**Fig. 1C**). Under InterPLM’s residue-level evaluation, a unit identifies one of 636 Swiss-Prot concepts when its thresholded activation matches the concept’s annotated residues with an F1 above 0.5 on both validation and test proteins(22). A random rotation of the same activations tests whether the neuron basis is privileged, because it keeps what the layer computes and changes only the axes(18).

We first planned this comparison over whole proteins, scoring a unit by whether it fires in the proteins that carry its term, and it separates nothing. Because most proteins lack a term and a unit is mostly silent, the median unit of each basis, including random rotations, tracks the best of its 100,228 candidate terms at a ROC AUC near 0.999 (**Supplementary Table 4**). We therefore judged the same units at the residues where curators located each concept, under prespecified rules. Average precision over the same proteins, a score that does not saturate, orders the units as the residue-level counts do (**Fig. 5D**).

**Fig. 5.**
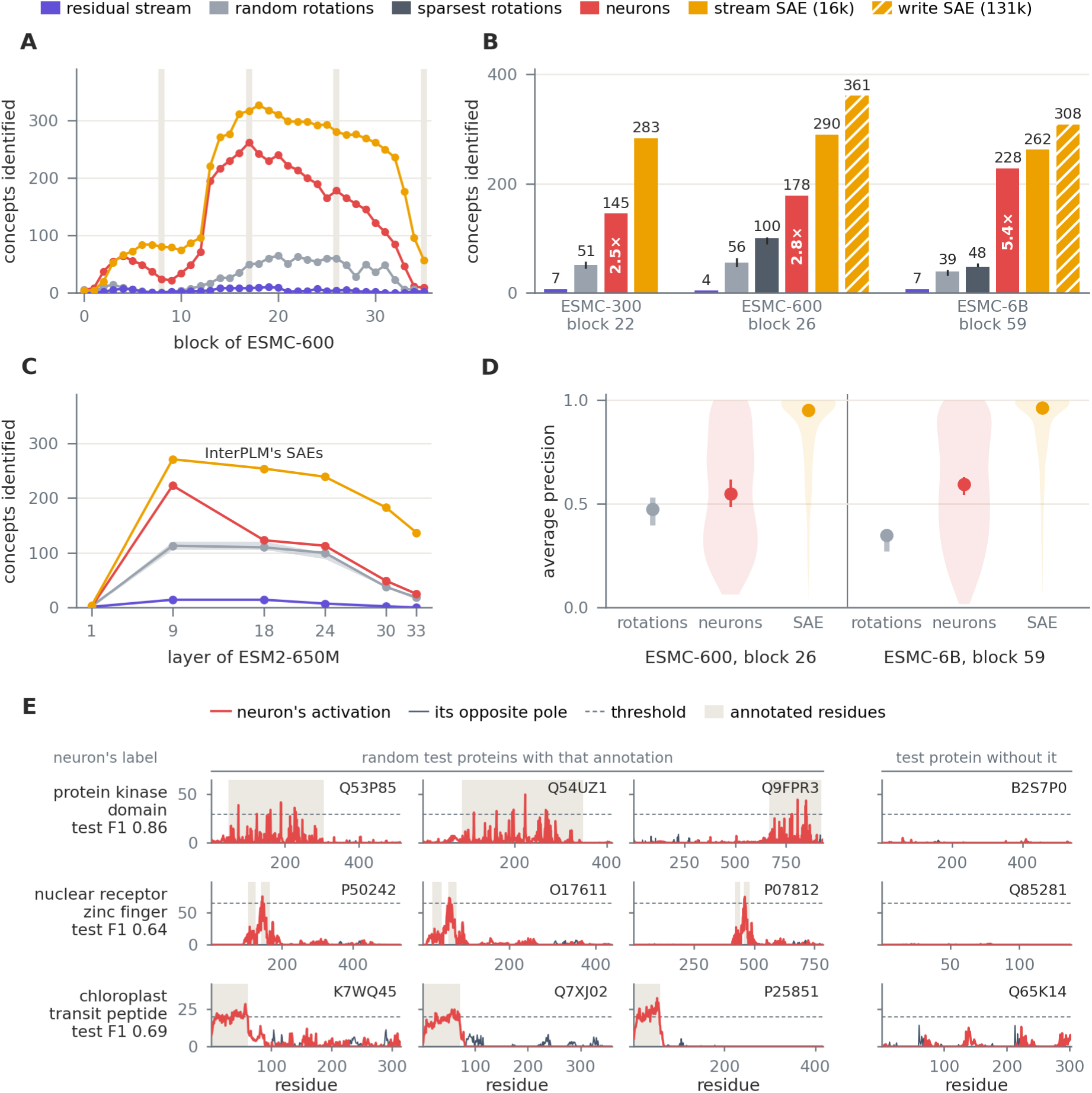
Read as poles, single feed-forward neurons identify at least 2.5 times as many residue-level concepts as random rotations of them, and fewer than SAEs. **A**, concepts, of 636, identified at every block of ESMC-600 by the stream’s dimensions, one random rotation, the neurons, and each block’s SAE on the stream (16,384 latents). Shaded columns mark the four blocks where the comparison was planned, and the neurons lead a rotation at 55 of the 56 blocks read in ESMC-600 and ESMC-6B. **B**, at three quarters of each model’s depth (blocks 22, 26, and 59), the depth at which Biohub released its SAE on the feed-forward write and its described SAE of ESMC-6B, the neurons identify 145 concepts in ESMC-300, 178 in ESMC-600, and 228 in ESMC-6B, which is 2.5, 2.8, and 5.4 times the best of ten random rotations. Also plotted are the sparsest rotations found by a search, Biohub’s SAE on the stream, and its SAE on the feed-forward write (131,072 latents). Bars show the median rotation for each copy, and whiskers span the range. **C**, under InterPLM’s own code in ESM2-650M, feed-forward poles against the residual-stream dimensions, three rotations (their median, with the range shaded), and InterPLM’s SAEs. **D**, over whole proteins, where the median unit of each basis tracks its best term at a held-out ROC AUC near 0.999, the held-out average precision of each unit’s best term at block 26 of ESMC-600 and block 59 of ESMC-6B. The neurons reach 0.55 and 0.59, the rotations reach at most 0.53 and 0.38, and the SAE reaches 0.95 and 0.96. Points are medians with 95% intervals from resampling units. **E**, three neurons from ESMC-600’s block 26, chosen after evaluating for recognizable biology, each on three test proteins drawn at random from those that carry its Swiss-Prot concept and on one test protein without it. Solid lines are the labeled pole’s activation along the residues and that of the same neuron’s opposite pole. Dashed lines mark the pole’s threshold and shaded residues mark those of the curated feature with the pole’s F1 on test proteins beside each name.

The neurons identify far more concepts than any rotation of them. At three quarters of each model’s depth, they identify 145, 178, and 228 concepts in ESMC-300, ESMC-600, and ESMC-6B, while no rotation identifies more than 57, 64, or 42 (**Fig. 5A,B**). Paired by concept, the neurons’ best test F1 exceeds that of each rotation (Wilcoxon signed-rank test, Holm-corrected *p <* 10*^−^*^55^). Under InterPLM’s own code, feed-forward poles of ESM2-650M identify 25 to 223 concepts a layer from layer 9 on, versus 0 to 14 for the residual-stream dimensions InterPLM compared(5) (**Fig. 5C**). Two signatures reported in vision and language models do not hold here, since rotations sparser than the neurons identify about half as many concepts in ESMC-600, and a neuron’s meaning does not narrow as it fires harder(32, 33) (**Supplementary Information, page 26**). SAEs still identify more, as each block’s SAE on the stream overtakes the neurons after the first few blocks and peaks at 327 concepts in ESMC-600 and 319 in ESMC-6B(43).

### Labels need held-out tests, and use appears in groups

Instead of having agents automate naming, we directly correlate model activations and annotations. Each pole is scored against every term of a curated annotation vocabulary, named by the term it tracks best on training proteins, and confirmed on held-out homology clusters(44) (**Fig. 1D** and **Fig. 5E**).

Despite leveraging poles, SAE dictionaries still add coverage that the neurons do not hold. At three quarters of each model’s depth, the neurons identify 179 of the 365 concepts that Biohub’s SAE on the feed-forward write identifies in ESMC-600 and 220 of 314 in ESMC-6B, and the SAE alone identifies most of the motifs, zinc fingers, and active sites that either identifies. The SAE’s latents are not the neurons found again, since fits of up to 64 poles track the latent for 1 of the 186 and 2 of the 94 concepts that only the SAE identifies (**Supplementary Fig. 5**).

The same scoring names a dictionary’s latents, and on Biohub’s described SAE of ESMC-6B the curated labels agree with the latents’ multi-agent descriptions above chance(24) (**Supplementary Fig. 5**). With annotation rows shuffled across proteins, however, 496 to 509 of a panel of 512 latents still received a label in each of 20 replicates. A label therefore counts here only when it is confirmed on held-out proteins (**Supplementary Information, page 28**).

A label on one neuron is not evidence of use, since setting a labeled pole of ESMC-600 to its resting value wherever it fires inside its term costs the masked prediction of those residues a median of only 0.0003 nats more than doing the same to placebo poles of matched activation (**Supplementary Fig. 6**). The poles that a concept’s label picks across blocks, a group assembled from labels, not discovered, are used together, since removing them at once lowers the log probability of the true residue at the concept’s masked tokens 37 times as far as placebo groups do and 23 times as far as the concept’s best single pole (**Fig. 4G**).

### Choosing where to read beats tuning what reads it

Finally, we asked whether knowing where a model is readable improves a downstream predictor of protein-protein interaction, a task on which the scores of published sequence-based predictors drop sharply once leakage between training and test proteins is removed(45). On the 2024 gold standard of Bernett and colleagues, along the stream every model reads interaction best in the first quarter of its depth, and a late feed-forward state of every model reads it better, by 0.02 to 0.06 MCC under our survey’s linear probes (**Fig. 6A,B**).

**Fig. 6.**
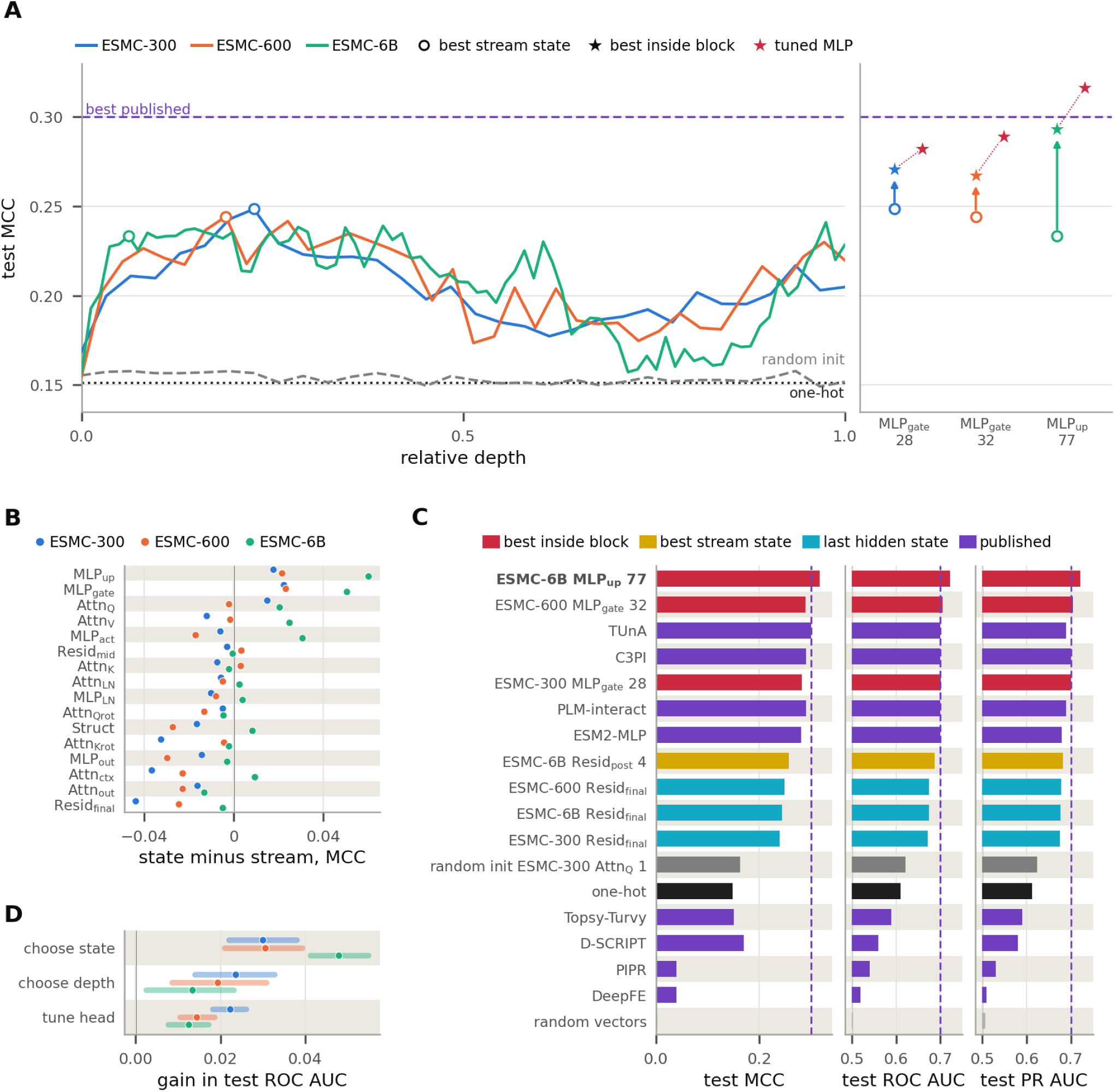
Where a frozen ESMC is read decides whether a simple probe reaches published protein-protein interaction predictors. All panels use the 2024 gold standard of Bernett and colleagues (52,048 test pairs). **A**, left, test MCC of our survey’s linear pair probe on each residual-stream state by relative depth, with each model’s best stream state (chosen on validation) as an open circle, beside the randomly initialized ESMC-300, the one-hot encoding, and the best published MCC. Right, for each model, the best stream state again, the linear probe at the best state inside a block (star, with the gain over the stream as an arrow), and a tuned one-hidden-layer MLP at that same state (second star). Only the tuned probe at ESMC-6B’s feed-forward up projection of block 77 passes the best published value. **B**, each state type’s best probe, the structure stream’s among them, minus the best residual-stream probe, both chosen on validation. **C**, test MCC, ROC AUC, and PR AUC of the tuned MLP at each model’s best state inside a block, at ESMC-6B’s best stream state, at each model’s last hidden state, and on the controls, beside published predictors (as reported to two decimals). Bars start at 0 for MCC, and at chance for the two areas, and dashed lines mark the best published value of each metric. **D**, gain in test ROC AUC under the same tuned MLP from reading the best state inside the block or the best stream state instead of the last hidden state, in addition to the gain from tuning the head on the last hidden state. Bars in C are means over three seeds, and intervals in D are 95% over bootstrap draws of the test proteins, paired between arms.

We refit the states our survey chose on validation with a tuned one-hidden-layer MLP. At ESMC-6B’s chosen state, the feed-forward up projection of block 77, this probe reaches an MCC of 0.316, a ROC AUC of 0.722 *±* 0.015, and a PR AUC of 0.721 *±* 0.017. The MCC is approximately 5% above the highest published value, 0.30, and both areas exceed the 0.70 or less reported by each recent predictor(46–48) (**Fig. 6C**). The same probe reaches an MCC of 0.257 on ESMC-6B’s best stream state and 0.24 to 0.25 on the last hidden state of every model. Choosing the state adds more than tuning the head does, 0.030 *±* 0.008 to 0.048 *±* 0.007 ROC AUC against 0.012 *±* 0.005 to 0.022 *±* 0.004 (**Fig. 6D**). A one-hot encoding of both partners reaches a ROC AUC of 0.610 *±* 0.016 under the same probe, so composition and length carry about half of what even the chosen state reads above chance, and the score is not evidence that the probe reads interfaces.

## Discussion

Checked against curated annotations, our conclusions about three ESMC models depended on where they were read, how a concept was decoded, which units were read, and how those units were scored. Scoring shows this most clearly: over whole proteins, a neuron, a random direction, and an SAE latent each track their best term almost perfectly, and only at annotated residues do the three separate. A whole-protein score is not wrong in itself, as average precision over the same proteins orders the units as the residue-level counts do, but a score near its ceiling cannot rank units. The reading that survives these checks needs no second trained model and no unit described by a language model. It complements SAEs and suggests an order of work in which a model is first read directly against curated labels and a dictionary is trained where the coverage that it adds is needed.

Single feed-forward neurons carry concepts that random directions through the same activity do not, in line with reports in vision and language models(32, 33, 49, 50). Our study adds a standard of judgment, since earlier work named and evaluated the unit roles with language models(30, 51), whereas we scored units against residues that curators annotated without reference to any model. The agreement is partial, as two signatures reported there do not appear in ESMC. The second pole adds 24% to 34% of concepts under SwiGLU and 2% to 12% under ESM2’s GELU. Dictionaries remain the more complete reading, since SAEs identify more concepts than the neurons at nearly every depth through latents that no group of up to 64 (topk) neurons tracks.

Between whole states and single neurons, the model’s use of what it holds appears at the scale of groups. A neuron can mark an active site where curators placed it, while removing it costs the model nothing, and the two results stop looking opposed once neurons are removed together, whether as a community found from the model’s own predictions or as the poles that share a concept’s label. The communities found here explain 2.6% of predictions, those the model is most sure of, not the curated concepts that single neurons mark(34).

For readouts, the directions a probe finds reflect the probe and its fitting as much as the states, since a per-label probe must fit each label from the few proteins that carry it(52). The common finding that larger pLMs do not improve linear-probe scores may therefore partly reflect the readout, and the state that a pLM returns by default is rarely the best place to read it(14, 53).

These failures are not particular to proteins: a score saturates wherever the population it scores over is mostly negative, and the unit is mostly silent; a probe’s lead reflects its fitting wherever training sets are small; and a label assigned without a held-out test is uninformative wherever the annotation is broad. Proteins provide the only large record of what a unit should respond to that no model wrote, and running these checks elsewhere needs an annotation that is located where a unit fires, curated independently of the model, and dense enough to score a unit where it is active.

The scope of each result limits its use, as our decomposition of the MLP’s lead, our interventions, and the communities ran on ESMC-600 alone. Erasure removes only what a linear map reads, and at the erased block a nonlinear probe still reads the concept, so the recovery we report is that of a linear readout(54). Curated annotation records only what curators have seen, so a unit without a label is not thereby a unit without a function.

Curated labels suggest a practice for anyone who reads a pLM. Choose a validation state, along the stream for most tasks, before taking the standard last hidden state as the model’s representation. Fit a linear probe to convergence before taking its failure as a concept’s absence. Label neurons as poles first, train a dictionary for concepts they miss, and test a labeled neuron’s use alongside neurons that share its label. Treat a concept’s readability, use, and recovery after erasure as three separate findings. We recommend future work analyzing if communities of neurons hold the concepts that only a dictionary identifies.

## Methods

### Models and states

We studied ESMC-300, ESMC-600, and ESMC-6B through their FastPLMs implementations(6, 23, 55, 56). ESMC models are bidirectional transformers trained with MLM, with 30, 36, and 80 blocks of width 960, 1,152, and 2,560 and SwiGLU feed-forward layers(42). Each block normalizes its input before attention and before the feed-forward network, normalizes queries and keys before the rotary position embedding, scales both writes to the residual stream before adding them, and applies a final layer norm after the last block(7, 36, 57). Forward hooks capture 17 types of state in one forward pass. They include the token embedding, 15 states from each block, and the output of the final layer norm, yielding 452, 542, and 1,202 states in the three models. The states of a block are the attention input after its layer norm, the queries, keys, and values, the queries and keys after rotation, the attention-weighted values, the attention output, the stream between attention and the feed-forward network, the feed-forward input after its layer norm, the gate and up projections, their product after the SiLU of the gate, the feed-forward output, and the stream after the block (**Supplementary Table 3**). The six state types that are split by attention head are also probed one head at a time. In the interaction and neuron analyses, we truncate proteins longer than 2,046 residues to their first 2,046.

ESMFold2 predicts structure from a frozen ESMC through one learned summary of its hidden states, *S* = Σ*_i_α_i_W_S_* LN(*X_i_*), which normalizes each stream state *X_i_* with a shared layer norm, projects it to 256 columns with a shared matrix *W_S_*, and sums the projections with learned softmax weights *α_i_*(23, 58). We probed *S*, the structure stream, as an additional state for each model, using the weights from that model’s ESMFold2 checkpoint. It brings the states read to 453, 543, and 1,203 in 18 types, and the best state of a model’s own state types excludes it.

Two of Protify’s baseline encodings serve as models whose only state is their token rows(59). A one-hot encoding of the residues pools to a protein’s composition and length, and a random-vector encoding, drawn from a seed fixed by the sequence, pools to its length alone.

### Datasets

We used the 22 concept datasets of Whitfield and colleagues with their published homology-aware splits, 18 from reviewed human entries of UniProtKB and InterPro and four from the BioMap benchmark, and the 2024 protein-protein interaction gold standard of Bernett and colleagues(14, 35, 37, 45, 60, 61). They cover 31 ProtParam values computed with Biopython, seven InterPro memberships, the three Gene Ontology aspects, localization, metal ion binding, and per-residue secondary structure, topology, peptides, functional sites, and modifications(62, 63). **Supplementary Table 7** lists each dataset with its source, size, and split.

### Controls and a randomly initialized model

We applied controls at test time to probes trained on the original inputs (**Supplementary Table 6**). They cross the model (trained, or an untrained copy of ESMC-300 with every module reset to its initialization), the probe (trained, or left at its initial weights), and the input (original, scrambled, unit-norm tokens, batch mean, or Gaussian), which gives 20 conditions for each full-width probe. The batch-mean control comes from single-cell perturbation studies, where a mean baseline often rivals complex models(64). Separately, we repeated the full probe analysis on a randomly initialized ESMC-300, with probes trained, selected, and scored on its own states, measuring what the architecture, the tokens, and a linear readout provide without any training. A randomly initialized transformer is not an empty baseline, since it can solve nontrivial tasks with only its embedding layers trained(65), and it acts as a control for sequence homology, because even under random weights similar sequence motifs attend to each other, so the states of similar sequences lie close together(59, 66).

### Probes

Protein-level probes read the mean and the variance of a state over a protein’s tokens, and residue-level probes read each residue’s state. Every token state is scaled to unit L2 norm before pooling or probing, a treatment that Whitfield and colleagues found to help residue probes and that we adopted by a rule fixed on ESMC-300 before any normalized result existed(14) (**Supplementary Information, page 18**).

A linear probe is *W* **h** + **b** on inputs standardized with the training split’s moments. We trained it with softmax cross entropy for multiclass tasks, per-label sigmoid cross entropy for multilabel and binary tasks, and squared error on standardized targets for ProtParam, using AdamW with early stopping on validation loss(41). One setting for each regime and loss, chosen on validation scores by sweeping learning rate, weight decay, and batch size, serves every probe and holds the fit fixed across the states that a map or a selection compares (**Supplementary Table 5**).

One AdamW setting with early stopping, the usual practice in probing studies of language models, cannot reach every probe’s optimum, so the linear probes at the states chosen on validation were also refitted to convergence, as transfer benchmarks in vision fit linear classifiers(40, 52, 67), with L-BFGS on the L2-penalized cross entropy for classification and ridge regression for ProtParam, the penalty chosen on validation(68) (**Supplementary Table 2**).

An MLP probe has one hidden layer, *W*_2_ GELU(*W*_1_**h** + **b**_1_) + **b**_2_, with dropout on the hidden units and the standardization, losses, optimizer, and early stopping of the linear probes(69). We swept its hidden width, dropout, learning rate, and weight decay on ESMC-300 validation scores. MLP and linear probes were compared on the 13 protein tasks at five state types of every model. To decompose the MLP’s lead, we trained the MLP with its activation replaced by the identity, a linear map of rank at most its hidden width that is trained like the MLP, and a reduced-rank regression with a penalty on both factors(70), on ESMC-600’s pooled means. Its difference from the MLP is the share of the lead that needs the nonlinearity, and its difference from the linear probe is the share that a hidden layer shared by a task’s labels gives.

The analyses differ in a few settings, each for a stated reason. The decomposition reads the mean alone, because the mean is linear in the token states where the variance is quadratic in them, so on the mean the MLP’s lead cannot come from second-order statistics that the input already supplies, and that lead is within 0.007 of the lead under the mean and variance (**Supplementary Information, page 22**). Cost sets the coverage of the other analyses. Linear probes read every state of all three models, while the decomposition, the interventions on the model’s objective, the communities, and the ablation of labeled poles ran on ESMC-600 alone, and the comparison of units at residues reads every block of ESMC-600 and every fourth block of ESMC-6B.

Classification probes are scored by the Matthews correlation coefficient (MCC), in Gorodkin’s multiclass form for multiclass tasks and as the binary MCC over all protein-label cells for multilabel and binary tasks(38, 71, 72). ProtParam probes are scored by the mean over test proteins of the cosine similarity between predicted and true standardized targets. Curves and maps show the test score of every state, with nothing selected. Where a result names the best state of a task, of a state type, or of a family of state types, that state is chosen on validation and reported on test, since the highest of hundreds of noisy test scores would overstate small test sets.

The output of block *k* of *n_b_* sits at a relative depth of (*k* + 1)*/*(*n_b_* + 1), and a task’s onset is the first relative depth at which its stream score comes within 5% of its highest. Tasks are grouped into six kinds of concepts, which are composition (ProtParam), secondary structure (three tasks), sorting and topology (localization, peptides, and topology), InterPro membership (seven tasks), function (the three Gene Ontology aspects and metal ion binding), and residue sites and modifications (functional sites, post-translational modifications, phosphorylation, and lipidation). A state type’s standing against the stream is its best probe over blocks minus the best stream probe of the same task, both chosen on validation.

A value written *x ± h* is a test score, or a difference of scores, with *h* half the width of its 95% interval over 1,000 bootstrap draws of the test proteins, and the same draws serve every model and probe of an analysis. ESMC-300’s survey, the MLP study, the tuned pair probes, the familiarity analysis, and the interventions on ESMC-600’s objective ran at three seeds and report means over them.

### Statistics

We report differences between probes, states, and models with 95% bootstrap intervals over the test proteins, paired between arms. Hypothesis tests are two-sided Wilcoxon signed-rank tests, paired by protein with a Bonferroni correction in the phosphomimetic scan, and paired by concept with Holm’s correction over the 23 rotations in the comparison of neurons with rotations, a test added after that comparison(73–75). SAE latent-term pairs are selected by the Benjamini-Hochberg procedure in one family over every latent and term, which also corrects the one-sided Fisher’s exact tests of curated terms at a community’s predictions, a test added after the community tests, and we test label agreement with descriptions by permutation(76, 77).

### Lenses, write ablations, and concept erasure

To test whether the concepts our probes read are used for the model’s own objective, we ran three analyses on ESMC-600 with its masked-language-model head. Each masks 15% of every test protein’s residues and scores the prediction of the masked tokens by accuracy, by the negative log-likelihood of the true residue, and by the KL divergence from the unaltered model’s prediction. The logit lens applies the final layer norm and head to the stream at every depth, and the tuned lens first applies an affine map learned for each depth(78, 79). Write ablations replace the attention or the feed-forward write of one block by its mean over training tokens.

Concept erasure uses LEACE, the least-squares concept eraser, fitted on the stream’s residue rows of 2,000 training proteins for each of four concepts (three-state secondary structure, localization, ProtParam composition, and InterPro family) and applied to every token after one of eight blocks while the model runs(39). The control erases as many random whitened directions in the same way. We scored the masked prediction of 1,000 test proteins and the concept’s linear readability at the erased depth, midway to the output, and at the final layer norm. Using criteria fixed before any analysis ran, we define a concept as used when erasing it costs the masked prediction more than the control (i.e., the difference is above zero), and as recovered when its readability at the final layer norm returns to at least half its unerased value. To separate what a linear erasure removes from what it leaves, we also erased two protein-level concepts from the mean-pooled stream at three blocks and refitted a linear probe, an MLP, and a linear probe on the variances over residues at the erased block and at later ones(54) (**Supplementary Information, page 21**).

### Familiarity

A model could score well on a protein that it recognizes from pretraining without encoding the concept, so we measured how familiar each test protein is to the model. The pseudo-log-likelihood (PLL) of a sequence of length *n* masks each residue in turn and averages the log probability of the true residue at the masked position, at the cost of *n* forward passes(80). We use a one-pass estimate, which we call the pseudo-pseudo-log-likelihood (PPLL), in which the unmasked sequence passes through the model once, and the log probability of each observed residue is averaged. Because the model sees each residue that it scores, PPLL is higher than PLL, but such one-pass scores are an established substitute for masked scoring under other names. Wild-type marginals from one unmasked pass predict variant effects nearly as well as masked scoring, a single-pass pseudo-perplexity estimates protein fitness, and a closed-form one-pass estimator of PLL correlates with it at a Spearman *ρ* of 0.92 in ESM2(81–83). PPLL is used here only to rank proteins by how predictable the model finds them.

For the 13 protein tasks we computed, task by task, Spearman’s *ρ* between PPLL and each encoder’s per-protein score at its best stream state, and the partial *ρ* between PPLL and ESMC-300’s gain over each control, controlling for length and label prevalence. By a criterion fixed before the run, a kind of concept reads as familiarity-driven if the median partial *ρ* over its tasks exceeds 0.1, with intervals that exclude zero on at least half of them.

### Protein-protein interaction

Protein-protein interaction is a task on protein pairs, based on the human 2024 gold standard of Bernett and colleagues, which shares no protein between splits and balances each protein’s positive and negative pairs(45). A pair probe reads the symmetric row [*z_a_* + *z_b_ | z_a_ ⊙ z_b_*] of the partners’ standardized pooled states *z_a_* and *z_b_*, which does not depend on the order of the partners, and is trained with sigmoid cross entropy. We report the MCC at a probability of 0.5, the ROC AUC, and the PR AUC, with intervals from bootstrap draws of the test proteins, so protein pairs move together.

For the tuned probes, we selected each model’s best state within a block and its best stream state, both chosen on validation under our survey’s linear probe, and then read them with a tuned linear probe and a tuned one-hidden-layer MLP. Each used the learning rate, weight decay, batch size, hidden width, and dropout that yielded the lowest validation loss, and we refit each with three seeds (**Supplementary Information, page 29**). We applied the same probes to one-hot encoding, a randomly initialized ESMC-300, and random vectors. Published values in **Fig. 6C** come from pair predictors that report all three metrics on this split, which are DeepFE, PIPR, D-SCRIPT, and Topsy-Turvy as re-evaluated by Bernett and colleagues, ESM2-MLP and TUnA from Ko and colleagues, and C3PI and PLM-interact from their own publications(9, 46–48, 84–87). PLM-interact’s MCC is implied by its reported precision and recall and is drawn at 0.29.

### Neuron communities

We discovered neuron communities in ESMC-600 after Shilov and colleagues(34). A prediction is the model’s output at one masked token of a protein with 15% of its residues masked. For each prediction, we score every feed-forward neuron by the absolute value of its activation times the gradient of the log probability of the model’s top residue with respect to that activation, and each block keeps its five highest. Over 541,883 predictions from 69,466 training proteins, two neurons are linked when they are kept together far more often than independent neurons would be, and the Leiden algorithm clusters the graph at eight resolutions into communities of 2 to 128 neurons(88).

We test each of 38,985 held-out predictions, from 2,500 proteins that share no cluster at 50% identity with a training protein, against the two communities that share the most neurons with the neurons kept for it, over spans of blocks. Within a span, a community explains a prediction when setting its neurons to zero moves the prediction by a KL divergence of at least 0.7 nats, keeping only its neurons moves it by at most 0.2, and setting every neuron of the span to zero moves it by at least 0.7, the thresholds of that work. The baseline reassigns each block’s neurons to the communities that hold them, preserving each community’s size within each block. Before the tests ran, the reading was that discovered communities explain a larger share of held-out predictions than shuffled ones, with the 95% interval of the difference above zero (**Supplementary Information, page 24**).

A concept’s group is the pole chosen for the concept in every ESMC-600 block where that pole’s test F1 is above 0.5. Every pole of the group is set to zero where it fires at the concept’s residues, and the fall in the log probability of the true residue at the concept’s masked tokens is compared with that of ten placebo groups, matched block by block for the activation removed, and with that of the group’s best single pole.

### Neuron poles

A feed-forward neuron of ESMC can push its output in either direction, and the two directions need not respond to the same thing, which is why Cammarata and colleagues read each neuron as two units(32). At each residue, a block’s feed-forward network forms **z** = SiLU(**g**) *⊙* **u** from the gate and up projections of its normalized input and writes *W_D_***z** to the stream(42). Each neuron thus adds a fixed vector to the stream in proportion to its activation, as in reading feed-forward layers as key-value memories(25). With the gate open, neuron *j* gives a positive pole 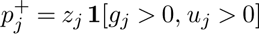 and a negative pole 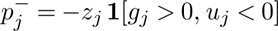. We omit the small outputs of a closed gate, as Tigges and colleagues also do(33). A rotation replaces **z** by *Q***z**, with *Q* a random orthogonal matrix, and splits each coordinate by sign. The residual stream’s dimensions are read both as InterPLM reads its neurons and splits by sign, and SAE latents, which are nonnegative, form one pole each(22).

The SAEs are Biohub’s(43). The figures draw the SAE of 16,384 latents on the stream leaving a block, the one size released for every block of all three models, and the SAE of 131,072 latents on the feed-forward write, released at ESMC-600’s block 26 and ESMC-6B’s block 59. These two blocks, and block 22 of ESMC-300, lie three quarters of the way through each model’s depth, where Biohub’s SAE on the feed-forward write and its described SAE of ESMC-6B read the model, so the comparisons at one depth are made there (**Supplementary Information, page 26**).

The residue-level comparison follows InterPLM’s evaluation with its code(22). Swiss-Prot proteins are labeled per residue for each concept, with half for validation and half for test; each unit’s activations are scaled by its maximum, and each concept’s best unit and threshold on validation are scored on test(35). A unit identifies a concept when its F1 exceeds 0.5 on both splits; the 20 concepts of amino-acid identity are left out, as in InterPLM’s report, leaving 636. We read ten rotations at three quarters of the depth of ESMC-600 and ESMC-6B, three in ESMC-300, and one at every block of ESMC-600 and every fourth block of ESMC-6B. In ESM2, at the layers of InterPLM’s released SAEs, the poles are the two signs of each neuron’s input to its GELU(5, 69). A GELU returns only a small value for a negative input, so the negative pole barely writes to the stream there, and the positive pole read alone identifies 219, 112, 103, 44, and 22 concepts at layers 9, 18, 24, 30, and 33, against 223, 123, 113, 49, and 25 for both signs. In ESMC, where SwiGLU lets both signs write, the positive pole alone identifies 107 of 145, 117 of 178, and 173 of 228 concepts at three quarters of the depth, so the second pole adds 24% to 34% of the concepts under SwiGLU and 2% to 12% under GELU.

To compare units over whole proteins, every unit at ESMC-600’s block 26 and ESMC-6B’s block 59 was labeled with an annotation compendium compiled from UniProtKB and its companion resources, split by cluster at 50% identity so that held-out proteins share no cluster with training proteins(35, 37, 44, 89, 90). Because SAE latents are sparse by construction and neuron poles are not, we first cut every unit to the SAE’s distribution of firing rates. A unit’s association with a protein-level term is the ROC AUC of its maximum over a protein for the term’s presence. We choose its best term on training proteins, then read that term’s ROC AUC and average precision on held-out proteins. We fixed the tests at this scale in advance: we wrote down and archived their metrics, thresholds, and decision rules before any activation was computed, and we decided them together after the run (**Supplementary Table 4**). We added the residue-level comparison afterward, with its rules written before it ran.

The sparsest-rotation search maximizes the sum of the fourth powers of the rotated, standardized activations by gradient ascent on the orthogonal group from three random starts(91). The dictionary test correlates each latent of the 131,072-latent SAE with every pole of its block over 2 million residues. A latent copies a pole when the two correlate at 0.7 or more, and groups of up to 64 poles are fitted greedily on one half of the residues and read on the other. To test whether the model uses what a labeled pole reads, each of 85 labeled poles of ESMC-600’s block 26 was set to its mean where it does not fire wherever it fires inside its term, and the fall in the masked tokens’ log probability was compared with that of ten placebo poles of matched activation. We selected the neurons drawn on proteins after evaluation, for recognizable biology, from the poles that are their concept’s best pole, and we drew each on test proteins picked at random with a fixed seed. They are not the block’s cleanest poles, since their test F1 of 0.86, 0.64, and 0.69 lies on either side of the median of 0.76 among the best poles of the 186 concepts whose test F1 exceeds 0.5.

### Use of large language models

The authors used Claude Opus 5.5 and Claude Fable 5.2 (Anthropic) to run automated scripts within the Protify framework that trained, scored, and summarized the probes; to write and run the code for the neuron study; and to assist in drafting this manuscript and its figures. The authors designed the study, directed and checked the analyses, and reviewed and edited the text, and take full responsibility for its content.

## DATA AND CODE AVAILABILITY

Every model, dataset and result of our study is gathered in the public Gleghorn Lab collection *Exhaustive hidden-state probing* on Hugging Face: the three ESMC models in their FastPLMs implementation, the 22 concept datasets with their homology splits, the annotation compendium that labels the neuron study’s atlas, and every score table, training record, selection, bootstrap interval and analysis table behind the figures(56). Those of the pooling and MLP study, the mean-only reruns, ESMC-600’s probe variants and state geometry, and its lenses, write ablations, and concept erasure are in a second archive. Probe weights and pooled stores are not archived, and every probe retrains from the released code and models. A third holds the neuron study’s record of the tests fixed in advance, atlas sample, per-unit scores, and evaluation tables.

The probing code is part of Protify, and the code that runs the neuron study and reproduces the figures and generated tables from the results, with the state-by-state maps left out of this paper, is in the plm_interpretability repository(59).

## AUTHOR CONTRIBUTIONS

Conceptualization: LH, JPG. Probing framework and software: LH. Formal analysis, investigation and data curation: LH. Writing, original draft: LH. Writing, review and editing: LH, JPG. Supervision: JPG. Funding acquisition: LH, JPG.

## ACKNOWLEDGEMENTS

This work was partly supported by a postdoctoral fellowship from the Institute for Engineering Driven Health at the University of Delaware and the National Science Foundation Accelerating Research Translation program 2331440 (LH), a University of Delaware Synthetic Biology Seed Grant (JPG), the National Institutes of Health through T32GM142603 (LH), and R01HL178817 (JPG), and the National Artificial Intelligence Research Resource Pilot Award (NAIRR240064). We thank Nikolaos Rafailidis and Tamar Peleg for their contributions to Protify, on which this study relies heavily, and for reviewing the work.

## Supplementary Information

The Supplementary Information follows the main text’s argument. Each of its first nine sections supports one section of the Results with controls, additional methods, and discussion that the main text has no room for, and none repeats a result shown in a main figure. The last section contains the tables the Methods point to. The main text and the Methods point to a section by its page.

### The depth map holds under controls, familiarity, and pooling

The depth map of **Fig. 2A** rests on linear probes of pooled states, so what it shows depends on how simply encodings read, whether a protein the model finds familiar scores more easily, and how states are pooled and normalized. Simple encodings read a large part of several concepts, so a probe’s score alone overstates what pretraining contributes (**Supplementary Fig. 1**). A one-hot encoding of the residues, which pooling turns into the protein’s composition and length, and a randomly initialized ESMC-300 both read composition nearly as well as the trained models. They also reach 0.26 to 0.69 on the InterPro tasks; for example, 0.61 and 0.63 on repeats, compared with 0.86 for the trained ESMC-300. Pretraining adds most where context decides the label, which residue identity alone cannot read, with a median gain over one-hot in ESMC-300 of 0.71 MCC for sorting and topology, 0.50 for secondary structure, and 0.40 for residue sites and modifications. Over the randomly initialized model, it adds a median of 0.44 for secondary structure, 0.41 for sorting and topology, 0.29 for residue sites and modifications, and 0.23 for InterPro membership, and nothing for composition. Lipidation is the exception, since residue identity reads it (0.24) better than any residual-stream state of any trained model (at most 0.13).

The randomly initialized ESMC-300, read by probes trained on its own states, measures what the architecture, the tokens, and a linear readout give without training. It reads much of the smallest InterPro tasks, 0.69 on binding sites and 0.51 on active sites against 0.81 and 0.65 for the trained model’s residual stream, and on the stream training adds 0.19 to 0.54 for families, domains, superfamilies, conserved sites, repeats, localization, topology, peptides, metal ion binding, functional sites, post-translational modifications, and secondary structure, 0.13 to 0.22 for the GO tasks, and all of phosphorylation (0.23), which the random model cannot read. Controls applied at test time stay near zero where the input carries no information about the label. At each task’s best stream state, the trained model’s probes applied to an untrained model’s states score within 0.05 of zero on every task except binding sites (*−*0.23) and repeats (*−*0.09), offsets that come from the score rather than from the states, since the micro-averaged MCC of a multilabel task penalizes a probe whose predicted label frequencies oppose the true ones. Scrambling the sequence preserves composition, and thus ProtParam (0.99), but leaves 0.16 for localization, 0.04 to 0.08 for secondary structure, and at most 0.03 elsewhere. The batch-mean control is not empty for a linear probe, whose output on the mean of eight states is the mean of its eight outputs, and it reaches 0.99 for ProtParam and 0.28 to 0.46 for localization, topology, metal ion binding, and post-translational modifications, where one label dominates a group of eight test proteins. Over the three seeds of ESMC-300’s survey, the standard deviation at each task’s best state is at most 0.031 and below the spread over test halves on 20 of the 22 tasks.

**Supplementary Fig. 1.**
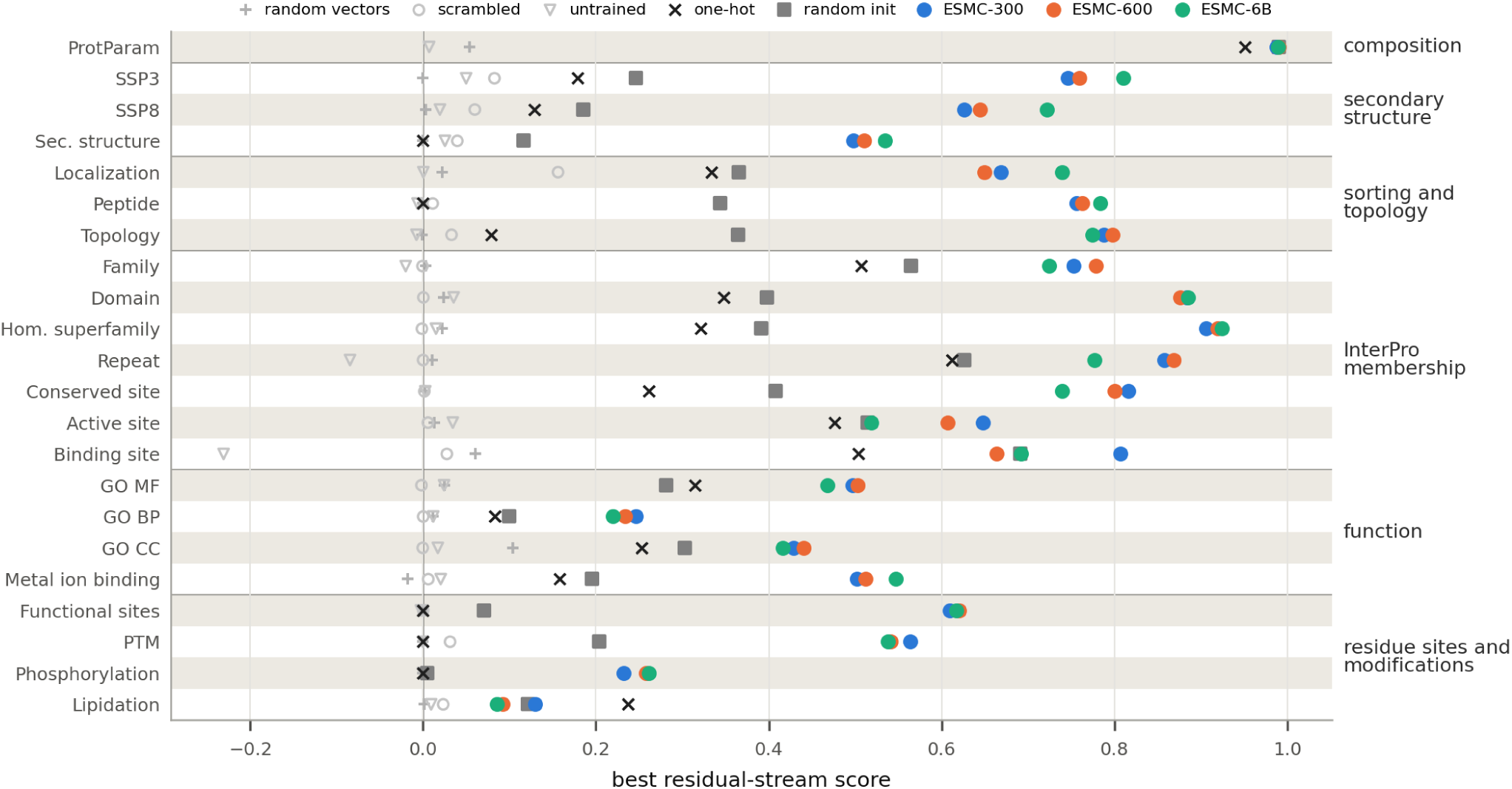
Pretraining adds most where sequence context decides the label. For each task, the best residual-stream score of each trained model (filled circles), of the randomly initialized ESMC-300 with its own probes, and of the one-hot and random-vector encodings. Hollow markers are ESMC-300’s probes at its best state applied to scrambled sequences and to an untrained model’s states. Scores are MCC and cosine similarity for ProtParam. The controls come close on composition and the small InterPro tasks, where pooled composition and length carry much of the label, and fall far behind on secondary structure, sorting and residue sites.

Part of what pretraining adds could be recognition rather than reading, since test proteins share little identity with the probes’ training proteins but they or their relatives were almost certainly in ESMC’s pretraining data, so a model could score well on proteins it recognizes without encoding the concept. We measured how familiar each test protein is to the model by its pseudo-pseudo-log-likelihood (PPLL), the mean log probability that one forward pass of the unmasked sequence assigns to each observed residue, which estimates the pseudo-log-likelihood in one pass(80) (**Methods, “Familiarity”**). PPLL ranks familiarity as intended, since ESMC-300 predicts calmodulin, histone H4, and ubiquitin, among the most conserved eukaryotic proteins, almost perfectly, at under 0.012 nats per residue against a median of 1.48. Yet probe correctness barely follows it, as Spearman’s *ρ* between PPLL and ESMC-300’s per-protein score lies between *−*0.49 and 0.23 over the 13 protein tasks (median 0.07, **Supplementary Table 1**), and the controls’ scores follow ESMC-300’s PPLL as much (medians 0.10 and 0.05). If pretraining’s lead came from recognition, the trained model’s gain over a control would rise with PPLL, which it does for no kind of concept. Controlling for length and label prevalence, the median partial *ρ* between PPLL and the gain over the randomly initialized model lies between *−*0.03 and 0.02 for every kind, and on InterPro families the gain falls with familiarity (*−*0.15 *±* 0.07), as the trained model’s lead over the seven InterPro tasks shrinks from 0.47 in the least predictable fifth of proteins to 0.35 in the most predictable. Pretraining’s lead therefore does not concentrate in the proteins a model finds predictable.

The protein probes read the mean and variance of a protein’s token states, on unit-norm states, and neither choice moves the map. Every trend holds when the survey is repeated on the mean alone, as the median onsets of ESMC-300 are unchanged, at 0.19 for residue sites and modifications, 0.23 for sorting and topology, 0.42 for InterPro membership, 0.45 for function, and 0.84 for secondary structure, the mean best stream score over the 22 tasks is 0.636, 0.625, and 0.629 against 0.634, 0.629, and 0.626, and the best of all states exceeds the best stream state by a median of 0.006, 0.008, and 0.012. On the 2024 gold standard, pair probes on the pooled mean stay within 0.013 ROC AUC of those on the mean and variance. Pooling the variance alongside the mean moves the mean-best score over the 13 protein tasks by at most 0.014 for either probe in any model, and where it costs linear probes, it costs them on the small InterPro tasks, where it doubles the width of probes trained on 482 to 2,097 proteins. Refitted to convergence, ESMC-300’s linear probes gain from the variance instead, by 0.008 *±* 0.005, so much of the cost comes from fitting twice as many inputs to fewer proteins. The one test fixed before the MLP study, that the variance helps MLP probes more than linear ones on ESMC-300, held by its rule (0.008 *±* 0.006), but the difference is small, absent in ESMC-600 (*−*0.001 *±* 0.006), and absent against converged linear fits (*−*0.004 *±* 0.006, 0.000, and 0.006 *±* 0.007 in the three models). Unit-norm token states, chosen on ESMC-300 by the rule of **Methods, “Probes”**, raise the mean score of the full-width probes from 0.501 to 0.535 over the 22 tasks but leave the best states nearly unchanged, with a mean best residual-stream score of 0.634 under both treatments, so normalization lifts weak probes far more than the best ones.

**Supplementary Table 1.**
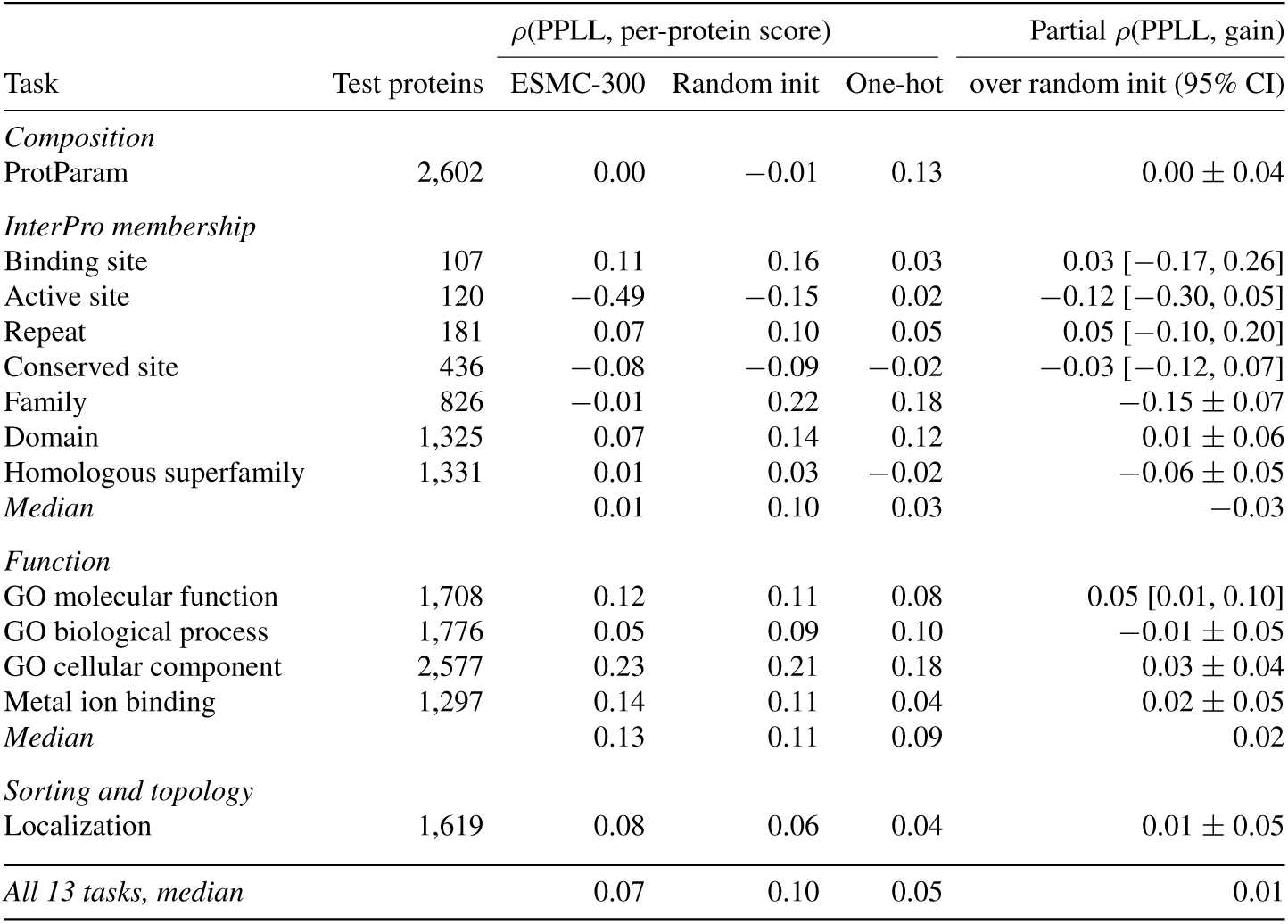
Probe correctness does not follow how predictable a protein is. For each of the 13 protein tasks, Spearman’s *ρ* between a test protein’s pseudo-pseudo-log-likelihood (PPLL) under ESMC-300 and its per-protein score, given for ESMC-300 and for the randomly initialized and one-hot controls, and the partial *ρ* between PPLL and ESMC-300’s gain over the randomly initialized model, controlling for length and label prevalence (**Methods, “Familiarity”**). Intervals are 95% over 1,000 bootstrap draws of test proteins. ESMC-300’s correlations have a median of 0.07 and the controls’ 0.10 and 0.05, so correctness follows familiarity no more for the trained model than for an untrained one. The criterion fixed before the run, that a kind of concept reads as familiarity-driven when the median partial *ρ* over its tasks exceeds 0.1 with intervals excluding zero on at least half of them, is met by no kind.

| Task | Test proteins | *ρ*(PPLL, per-protein score) |  |  | Partial *ρ*(PPLL, gain) |
|---|---|---|---|---|---|
|  |  | ESMC-300 | Random init | One-hot | over random init (95% CI) |
| *Composition* |  |  |  |  |  |
| ProtParam | 2,602 | 0.00 | −0.01 | 0.13 | 0.00 ± 0.04 |
| *InterPro membership* |  |  |  |  |  |
| Binding site | 107 | 0.11 | 0.16 | 0.03 | 0.03 [−0.17, 0.26] |
| Active site | 120 | −0.49 | −0.15 | 0.02 | −0.12 [−0.30, 0.05] |
| Repeat | 181 | 0.07 | 0.10 | 0.05 | 0.05 [−0.10, 0.20] |
| Conserved site | 436 | −0.08 | −0.09 | −0.02 | −0.03 [−0.12, 0.07] |
| Family | 826 | −0.01 | 0.22 | 0.18 | −0.15 ± 0.07 |
| Domain | 1,325 | 0.07 | 0.14 | 0.12 | 0.01 ± 0.06 |
| Homologous superfamily | 1,331 | 0.01 | 0.03 | −0.02 | −0.06 ± 0.05 |
| *Median* |  | 0.01 | 0.10 | 0.03 | −0.03 |
| *Function* |  |  |  |  |  |
| GO molecular function | 1,708 | 0.12 | 0.11 | 0.08 | 0.05 [0.01, 0.10] |
| GO biological process | 1,776 | 0.05 | 0.09 | 0.10 | −0.01 ± 0.05 |
| GO cellular component | 2,577 | 0.23 | 0.21 | 0.18 | 0.03 ± 0.04 |
| Metal ion binding | 1,297 | 0.14 | 0.11 | 0.04 | 0.02 ± 0.05 |
| *Median* |  | 0.13 | 0.11 | 0.09 | 0.02 |
| *Sorting and topology* |  |  |  |  |  |
| Localization | 1,619 | 0.08 | 0.06 | 0.04 | 0.01 ± 0.05 |
| *All 13 tasks, median* |  | 0.07 | 0.10 | 0.05 | 0.01 |

The onsets of **Fig. 2A** do not rest on single noisy depths, since on a running mean over three depths 61 of the 66 onsets move by at most 0.05 and no kind’s median onset in any model moves by more than 0.04, with the order of the kinds unchanged except where two medians lay within 0.02 of each other (residue sites and modifications against sorting and topology in ESMC-600). A stricter rule, the first depth from which a task’s score stays within 5% of its highest for three depths in a row, leaves composition at the embedding, secondary structure last (0.84, 0.89, and 0.94), and InterPro membership earlier in ESMC-6B than in the smaller models (0.37 against 0.42 and 0.41), but it moves 17 onsets by more than 0.05 and finds no such run for 9 tasks, among them lipidation in every model, whose scores stay near that line over much of the depth, so one task’s onset is less certain than its kind’s place in the order.

Principal components show the same states that need no probe (**Supplementary Fig. 2**). For five protein tasks we projected each protein’s pooled mean state after blocks 0, 5, 10, 15, 20, 25, and 29 of ESMC-300, on raw states at one seed, onto its first two principal components, computed over the task’s proteins, and projected scrambled copies of the test proteins onto the same components. We used principal components rather than nonlinear embeddings, whose results depend strongly on their settings(92). InterPro families group in the trained model, most clearly in the middle blocks. Measured as the share of a protein’s ten nearest neighbors in the projection that belong to its own family set, grouping rises for rhodopsin-like G protein-coupled receptors from 0.31 at block 0 to 0.90 at block 10 and reaches 0.99 for immunoglobulin light chains and 0.68 for S1A peptidases at block 20, and the sets mix again at block 29 (0.08 to 0.63). Scrambled sequences separate from natural proteins from block 15 (0.93). The randomly initialized model keeps the neighborhoods of its embedding at every block (0.05 to 0.69), and its first component explains about 20% of the variance at every block, against 22 to 64% in the trained model.

**Supplementary Fig. 2.**
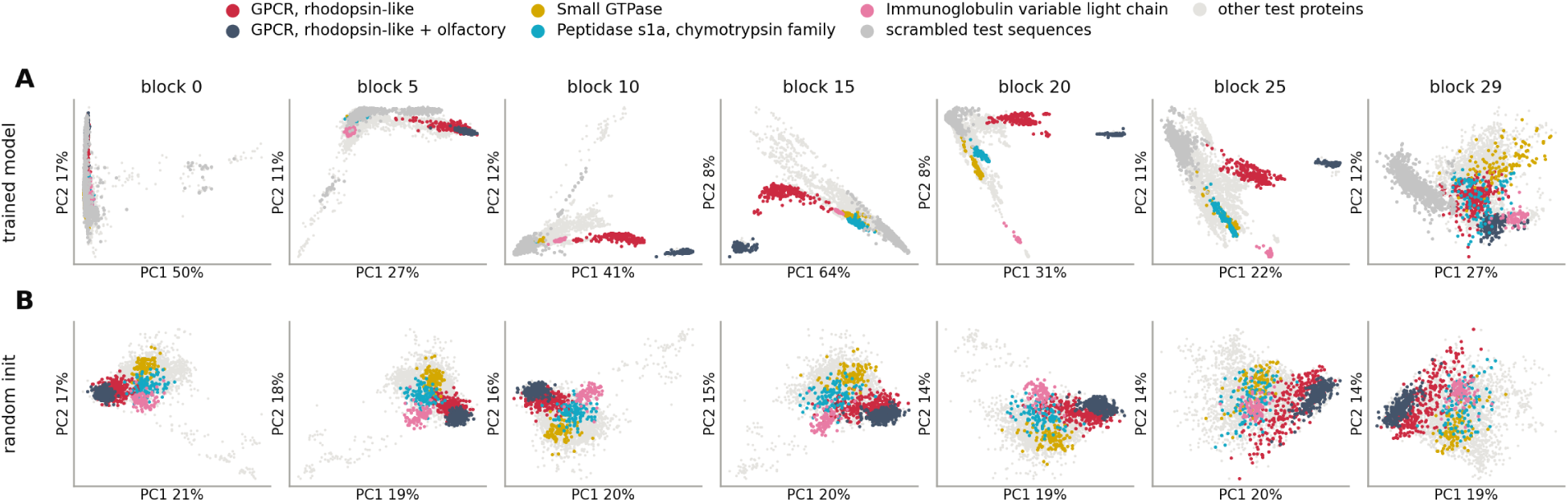
Principal components of pooled states. Each protein’s pooled mean state after blocks 0 to 29 of ESMC-300 (columns), projected onto its first two principal components, for the trained model (**A**) and the randomly initialized model (**B**), with the five most frequent InterPro family sets and the scrambled copies of the test proteins marked as in the key. Scrambled copies are projected onto the trained model’s components. Families group most clearly in the middle blocks of the trained model and mix again at the last block, while the randomly initialized model keeps the neighborhoods of its embedding throughout.

### A concept is decodable long before it is needed, and is linearly readable again after erasure

Erasing a concept early costs the masked prediction almost nothing (**Fig. 2B**), and lenses and write ablations show why, by locating where ESMC-600’s stream holds the prediction and where the writes that the objective needs begin. The prediction becomes readable in the last fifth of the model’s depth (**Supplementary Fig. 3A**). Read through the model’s own final norm and head, the logit lens, the stream predicts masked tokens no better than always guessing the most common residue (accuracy 0.086) up to a relative depth of 0.78(78). Accuracy then rises to the model’s final 0.544 over the last seven blocks. At mid-depth it even falls below that baseline, to 0.001 at relative depths 0.51 to 0.54, where the loss of 6.1 nats per masked token is worse than a uniform guess over the vocabulary (3.5 nats), so that read directly by the head the stream there is confidently wrong. A tuned lens, which learns an affine map for each depth, reads more from early states but still reaches half of its total gain only at 0.83(79). Each unmasked token’s own identity stays decodable at every depth (accuracy at least 0.997), yet the loss of this identity readout climbs from under 0.002 nats in the first third of the depth to a peak of 0.036 at a relative depth of 0.68 and falls back by the last blocks (**Supplementary Fig. 3B**). The peak sits where pooled composition, a count of token identities, is least readable.

Write ablations, which replace one block’s attention or feed-forward write by its mean, locate the computation the objective needs (**Supplementary Fig. 3C**), as a single replacement costs almost nothing in the first 13 of ESMC-600’s 36 blocks, a median of 0.004 nats per masked token against a clean loss of 1.46, except for the attention write of block 0 (0.07). Feed-forward writes become necessary abruptly at block 13 (relative depth 0.38), costing 0.06 to 0.35 nats, and attention writes follow at block 19 (0.54), where the logit lens collapses. The costs peak around a relative depth of 0.65 and again in the last blocks. Each early block is thus individually redundant for the objective, while every middle and late block contributes.

**Fig. 2B** omits the first erased depth because, after the first block, the random control costs the masked prediction more than LEACE for every protein-level concept, since random whitened directions there remove token identity(39). Every concept is used at every depth from a relative depth of 0.57 on. Readability at the final layer norm after erasure recovered to at least half of its unerased value at every depth up to 0.89, and failed to recover only in the last block, for secondary structure and composition, so a concept erased anywhere before the last block is linearly readable again at the output, and readability at the output says little about where a concept is computed. Fel and colleagues describe one route to such recovery: a linear erasure equalizes the classes’ means and leaves the concept in their covariances, where a nonlinear probe still reads it and later nonlinear layers return it to a linear code(54). We tested this account on two protein-level concepts, localization and metal ion binding, by erasing each from the mean-pooled stream with LEACE at blocks 8, 17, and 26. At the erased block, a refitted linear probe drops to a balanced accuracy of 0.32 to 0.38, from 0.65 to 0.67 on the intact model, while a refitted MLP keeps 0.59 to 0.61 and a linear probe on the variances over residues 0.59 to 0.63. Two blocks later, the refitted linear probe has regained 72 to 89% of its intact accuracy above the majority class. The concept therefore stays at the erased block for a nonlinear reader, and the later blocks, which are frozen here where the model of Fel and colleagues was trained with the eraser in place, return it to a linear code. Of the other tests that we fixed before any of these analyses ran (**Methods, “Lenses, write ablations, and concept erasure”**), the identity readout fell mid-depth as set, and the logit lens stayed within 0.05 of the most common residue’s frequency only from above, since accuracy falls below it mid-depth, so that test failed as written.

**Supplementary Fig. 3.**
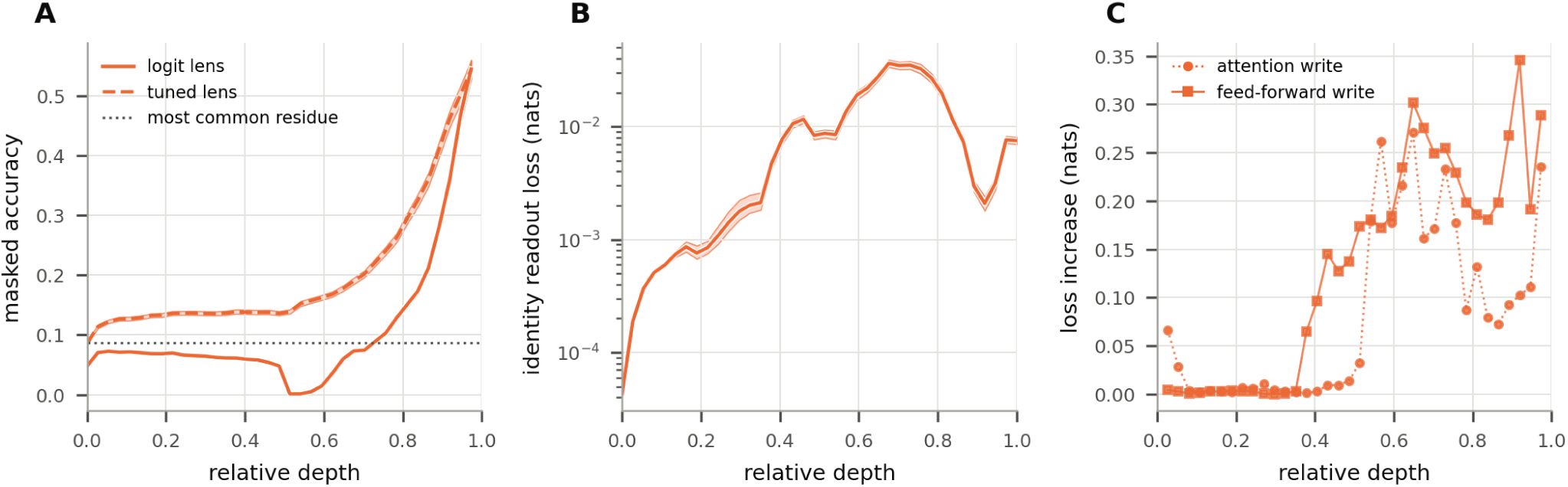
ESMC-600’s masked prediction becomes readable only in the last fifth of its depth, though the writes it needs start near a relative depth of 0.4. All panels use the test proteins of the three-state secondary-structure task with 15% of residues masked. **A**, masked-token accuracy of the stream at each depth, read through the model’s own final norm and head (logit lens) or through an affine map learned per depth first (tuned lens). The dotted line is always predicting the most common residue. **B**, loss of a linear readout of each unmasked token’s own amino acid (lower is more explicit). Identity is least explicit near a relative depth of 0.7, where pooled composition dips. **C**, rise in masked-token loss when one block’s attention or feed-forward write is replaced by its mean, at the block’s depth. Bands are 95% intervals over test proteins. Early blocks are individually redundant, while feed-forward writes matter from a relative depth of about 0.4 and attention writes from about 0.55.

### Fitting and the shape of the readout explain most of the MLP’s lead

The main text attributes most of the MLP’s lead to fitting and to the form of the readout (**Fig. 2C,**D), and the evidence for each lies task by task. One global AdamW setting with early stopping understates what a linear readout can reach(41) (**Supplementary Table 2**). Refitted with converged, validation-tuned logistic and ridge regressions, the best probes of the 13 protein tasks of ESMC-300 gained a median of 0.019 at the best state of any type and 0.049 at the best stream state, up to 0.11 for InterPro binding sites, the smallest training set (446 proteins). Refitted, the best stream state matched or beat the refitted best state of any type on 7 of 13 protein tasks, so part of the edge that states inside a block hold under AdamW is fitting, and comparisons between states and models share the AdamW bias, which is largest on small training sets. In the MLP study, converged fits of the protein tasks of all three models beat the AdamW probes by 0.038 to 0.040 on average at the best state of any type.

**Supplementary Table 2.** Converged solvers against the global AdamW setting. For each protein task of ESMC-300 (one seed, unit-norm states), the full-width probe chosen on validation among all state types and among residual-stream states (all Resid_post_, given by block), refitted with a validation-tuned L2 penalty by L-BFGS (ridge regression for ProtParam)(68). Scores are test MCC, and cosine similarity for ProtParam. Converged fitting lifts the stream’s probes more than those of the best state of any type, so part of the intra-block states’ edge under AdamW is fitting.

|  | Best state of any type |  |  |  | Best residual-stream state |  |  |  |
|---|---|---|---|---|---|---|---|---|
| Task | State | AdamW | Solver | Change | Block | AdamW | Solver | Change |
| ProtParam | Resid<sub>mid</sub> 1 | 0.989 | 0.991 | +0.002 | 1 | 0.988 | 0.990 | +0.002 |
| Localization | Attn<sub>ctx</sub> 28 | 0.678 | 0.728 | +0.050 | 6 | 0.667 | 0.718 | +0.051 |
| Family | MLP<sub>gate</sub> 28 | 0.820 | 0.809 | −0.011 | 18 | 0.752 | 0.822 | +0.070 |
| Domain | Resid<sub>mid</sub> 24 | 0.890 | 0.891 | +0.001 | 24 | 0.885 | 0.889 | +0.005 |
| Homologous superfamily | Attn<sub>Q</sub> 27 | 0.921 | 0.906 | −0.015 | 26 | 0.902 | 0.912 | +0.010 |
| Repeat | Attn<sub>K</sub> 12 | 0.880 | 0.892 | +0.011 | 5 | 0.854 | 0.887 | +0.033 |
| Conserved site | Attn<sub>K</sub> 16 | 0.829 | 0.843 | +0.013 | 28 | 0.811 | 0.847 | +0.036 |
| Active site | Attn<sub>K</sub> 8 | 0.668 | 0.707 | +0.039 | 29 | 0.649 | 0.739 | +0.090 |
| Binding site | Attn<sub>Q</sub> 6 | 0.835 | 0.926 | +0.091 | 5 | 0.807 | 0.921 | +0.114 |
| GO MF | Attn<sub>K</sub> 24 | 0.513 | 0.532 | +0.019 | 25 | 0.494 | 0.547 | +0.053 |
| GO BP | Attn<sub>K</sub> 24 | 0.243 | 0.275 | +0.032 | 23 | 0.234 | 0.283 | +0.049 |
| GO CC | Attn<sub>Q</sub> 27 | 0.435 | 0.475 | +0.040 | 7 | 0.445 | 0.465 | +0.019 |
| Metal ion binding | Attn<sub>LN</sub> 20 | 0.480 | 0.549 | +0.069 | 22 | 0.494 | 0.560 | +0.066 |

The MLP study read five state types (Resid_embed_, Resid_post_, Resid_final_, Attn_ctx_, MLP_act_) and chose each task’s best state among them, and its linear probes, on stores written anew, match the survey’s probe by probe to a median absolute difference of 0.001 at the seed both share. The MLP’s lead is largest on the small InterPro tasks, binding and active sites, at 0.15 to 0.31 over AdamW in every model and, on ESMC-300, 0.035 and 0.167 over converged fits, and absent on domains and homologous superfamilies. It grows with scale because the two probes move apart, as the AdamW linear probes fall from 0.696 to 0.684 and 0.670 while the mean best MLP score holds at 0.757, 0.758, and 0.767. Ranking metrics agree with MCC, since on the 12 classification tasks macro ROC AUC is 0.918 to 0.926 for MLP probes across the three models against 0.860 to 0.868 for linear probes, average precision 0.682 to 0.689 against 0.612 to 0.655, and balanced MCC 0.837 to 0.844 against 0.754 to 0.770. On the randomly initialized ESMC-300, MLP probes add 0.013 *±* 0.008 over AdamW linear probes, against 0.061 on the trained model.

The mid-depth dip of InterPro membership that linear probes show in every model all but disappears under MLP probes, as **Fig. 2D** shows for ESMC-600. Over the InterPro tasks, MLPs lose at most 0.03, 0.02, and 0.04 of their score at the middle depths of ESMC-300, ESMC-600, and ESMC-6B, against up to 0.36, 0.26, and 0.38 for linear probes, and in ESMC-6B, where the linear dip is deepest, conserved sites fall from 0.76 to 0.39 under linear probes and from 0.86 to 0.82 under MLPs. Composition behaves differently, since ProtParam’s dip persists under MLP probes in every model (to 0.80, 0.79, and 0.78, against 0.76 for linear probes in all three), so composition, unlike membership, is less readable mid-depth. In the last blocks the order reverses, and at the final layer norm MLPs read InterPro conserved sites, domains, families, and homologous superfamilies 0.07 to 0.17 lower than linear probes in every model.

Five probes separate what the hidden layer adds from what its activation adds, on ESMC-600’s unit-norm pooled means, which are the decomposition’s design and on which the MLP’s lead over converged linear fits is the same within 0.007 on average as under the main text’s mean and variance. Beside the MLP without its activation (**Methods, “Probes”**), we trained a quadratic probe, a linear probe on the mean and its elementwise square under the linear probes’ settings, the smallest step beyond a linear readout. At each task’s best state of any type, the mean score over the 13 tasks is 0.685 for the AdamW linear probe, 0.674 for the quadratic probe, 0.725 for the converged linear fit, 0.748 for the MLP without its activation, and 0.760 for the MLP. The activation therefore adds 0.012 *±* 0.004 of the MLP’s 0.075 lead over the AdamW linear probe, and the MLP without its activation beats the converged fit by 0.023 *±* 0.008. By kind, the five probes score 0.789, 0.772, 0.831, 0.878, and 0.890 on InterPro membership, 0.421, 0.420, 0.465, 0.460, and 0.475 on function, 0.702, 0.690, 0.758, 0.744, and 0.752 on localization, and 0.991 to 0.993 on composition. Task by task, the activation matters most on InterPro conserved sites (0.068 *±* 0.024), where the MLP without its activation scores below even the AdamW linear probe, and on Gene Ontology molecular function (0.029), and not at all on repeats, where the MLP without its activation scores 0.017 higher. The MLP without its activation gains most over converged fits on active sites (0.306 *±* 0.070) and repeats (0.061), and trails them on binding sites (*−*0.036), homologous superfamilies (*−*0.016), and Gene Ontology cellular component and molecular function (*−*0.009 and *−*0.011). The split is alike at each task’s best stream state, where the MLP leads the linear probe by 0.085, the MLP without its activation by 0.065, and the activation adds 0.019 *±* 0.004. The squared terms cost the linear probe 0.010 *±* 0.008 on average, most on families (*−*0.093) and repeats (*−*0.034), whose probes they double in width on 2,097 and 819 training proteins. The criterion fixed in advance, that an MLP without its activation within 0.02 of the linear probe would leave the gap to the nonlinearity, fails, because the hidden layer’s shape and training carry most of the gap.

### Standardized inputs let an untrained model pass an intervention test

How a readout’s input is treated can decide an intervention test as well as a probe score. We repeated the phosphomimetic scan of Whitfield and colleagues on ESMC-300, with 1,000 phosphoproteins drawn at random from those with annotated phosphoserine, phosphothreonine, or phosphotyrosine sites(14). Every serine, threonine, and tyrosine was mutated to alanine, glycine, and glutamate. A mutant is labeled active when it keeps an annotated site’s function (the wild type, or glutamate at an annotated phosphosite), inactive when it removes it (alanine or glycine at an annotated phosphosite), and unknown at any other serine, threonine, or tyrosine. For every state type and block, a three-class logistic regression on the pooled mean state, fitted with scikit-learn on a stratified 70:30 split of the protein’s mutants, predicts the label(93). Controls are the same regression on shuffled labels and on the states of the untrained model, a copy of ESMC-300 with every module reset to its initialization. Each regression is fitted on raw means, as in the original protocol(94), and on means standardized over the protein’s training mutants. Each protein’s MCC is averaged over three replicates before a Wilcoxon signed-rank test of true against shuffled labels, paired by protein, with a Bonferroni correction over the 452 states of each input treatment(73, 74).

The test reproduces on ESMC-300 (**Supplementary Fig. 4**). On raw pooled means, the trained model separates mutants that keep a phosphosite’s function from those that remove it only in the last blocks, reaching an MCC of 0.35 at block 29, while shuffled labels and the untrained model stay at zero, and 203 of 452 states pass the corrected test. Standardizing the means over each protein’s training mutants changes the picture, since the trained model then reaches 0.28 from block 0 and 0.40 by block 28, and 447 of 452 states pass, but the untrained model reaches 0.21 as well. One substitution changes one of hundreds of tokens and moves a pooled mean very little, and standardizing amplifies the substitution itself, which any network encodes. Only the trained model’s margin over the untrained one, a median of 0.15 along the stream, is therefore attributable to training.

**Supplementary Fig. 4.**
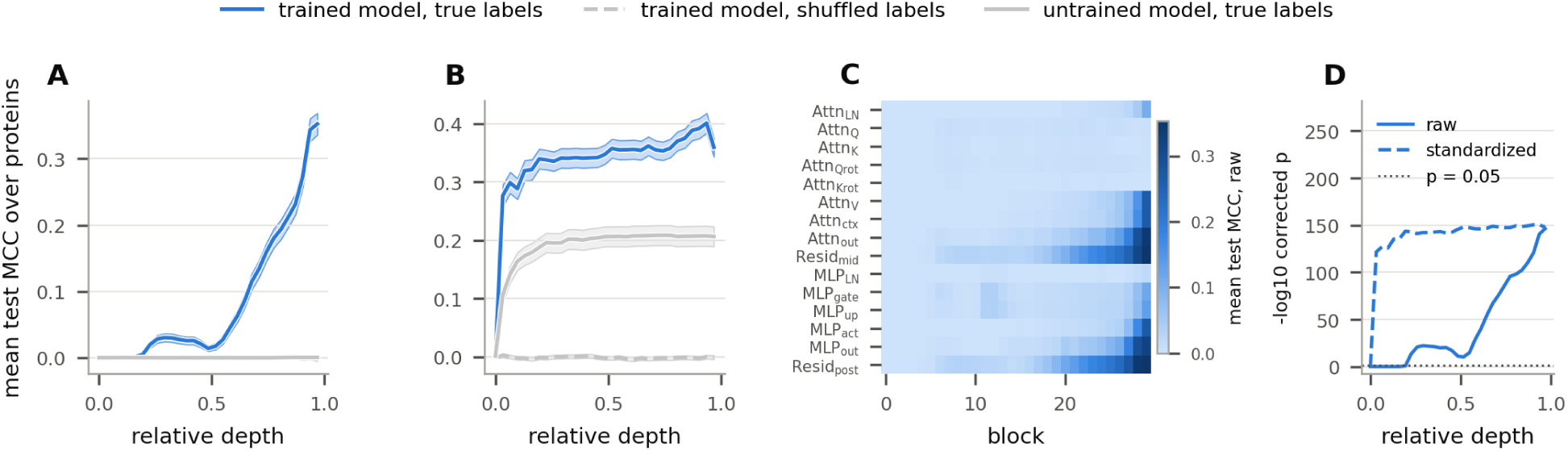
Phosphomimetic scanning in ESMC-300. Logistic regressions on pooled mean states of mutants of 1,000 phosphoproteins predict whether a mutant keeps an annotated phosphosite’s function, removes it, or is unrelated. **A** and **B**, mean test MCC over proteins along the residual stream on raw (**A**) and standardized (**B**) means, for the trained model with true and with shuffled labels and for the untrained model, with 95% intervals shaded. **C**, the trained model’s MCC on raw means at every state type and block. **D**, the Bonferroni-corrected *p* value of the Wilcoxon test of true against shuffled labels along the stream on raw and standardized means, with the dotted line at 0.05. On raw means only the last blocks separate the classes, and standardizing lets even the untrained model separate them, so only the trained model’s margin over it reflects training.

### Inside a block, the large gains fall on the smallest tasks

The main text groups the tapped state types by how far they lie from the stream (**Fig. 3A** and **Supplementary Table 3**), and task by task the same pattern holds in every model. The branch inputs and the stream between the branches match the stream, the rotated queries and keys and the SwiGLU product fall short across tasks, the final layer norm falls short on residue tasks, and the large positive differences sit on the small InterPro tasks and lipidation, where training sets are smallest. In ESMC-300, the attention input of block 0 reads lipidation at 0.24 against 0.13 for the embedding, the best stream state, and the keys of block 7 read binding sites at 0.89 against 0.81. In ESMC-600, the feed-forward gate of block 0 reads lipidation 0.28 above the stream. In ESMC-6B, mid-depth queries, keys, and feed-forward states read binding, active, and conserved sites and repeats 0.06 to 0.27 above the stream, the feed-forward gate of block 18 reads lipidation 0.53 above it, and the other residue tasks gain at most 0.05 inside the block. Because converged fitting lifts the stream’s probes more than those of the best state of any type (**Supplementary Table 2**), part of these gains is fitting. The structure stream, the last column of **Fig. 3A**, keeps InterPro membership and function and reads composition and residue sites and modifications 0.15 to 0.23 below the stream.

The lead of attention’s projections on binding and active sites (**Fig. 3C,D**) belongs to the pooled readout and does not show tokens that the stream lacks. Those two tasks are InterPro memberships read from a protein’s pooled states, whereas on functional sites, a residue-level task whose probes read unpooled tokens, the stream’s chosen state reads at least as well as the chosen query or key state in every model (0.610, 0.620, and 0.617 against 0.594, 0.605, and 0.605 in ESMC-300, ESMC-600, and ESMC-6B). Under pooling, the projections lead in every model, by 0.09, 0.08, and 0.21 on binding sites and by 0.03, 0.04, and 0.27 on active sites. The two groups of tasks take their labels from different sources, so this comparison is consistent with pooling as the cause without isolating it.

Single attention heads were probed as well, and in ESMC-300 the best single head, 64 dimensions chosen among 2,700 head probes, comes within 0.06 of the best full-width probe on 12 of the 13 protein tasks, while on residue tasks it falls 0.06 to 0.24 short on 7 of 9. On four InterPro tasks the best head beats the best full-width probe, by 0.17 on active sites, 0.05 on conserved sites, and 0.01 on repeats and binding sites, which, with 128 pooled features instead of 1,920, points to the size of the probe rather than to a specialized head. For protein tasks, depth separates heads far more than identity does, as the variance of head scores between blocks is 6 to 57 times the variance among the heads of one block, with metal ion binding (1.4) the exception, whereas for residue tasks the ratio is 0.12 to 4.9, so individual heads differ about as much as blocks do. Rotary position embedding lowers what queries and keys carry for every task but peptides, by 0.045 for queries and 0.042 for keys averaged over blocks and tasks, while peptides gain 0.015 to 0.033, because their signal and transit peptides sit at the N terminus and rotation writes absolute position into the state.

### Communities and groups of neurons: discovery, tests, and limits

**Fig. 4** reports communities of neurons in ESMC-600 by a reading fixed before the tests ran (**Methods, “Neuron communities”**), and how the tests were run bounds the share of predictions that they explain. The attribution keeps neurons whose gate is open, since 19.8% of the cells that it keeps have a closed gate, against 74.0% of all cells at predictions. Of the model’s 110,592 feed-forward neurons, 107,668 are kept at some prediction, and the 27 that are kept at more than 20% of predictions were set aside as always on. The graph has 10.9 million links, and its 39,260 communities are small, with a median of 2 neurons in 2 blocks and a 90th percentile of 6 neurons in 5 blocks. A discovered community explains 2.62% of held-out predictions within a span of at least 1 block, 2.21% within at least 4, and 0.74% within at least 12, against 0.010%, 0.003%, and 0 for shuffled communities, with differences of 2.61% [2.45, 2.77], 2.21% [2.06, 2.36], and 0.74% [0.65, 0.83]. This is about a quarter of the coverage that Shilov and colleagues report for a language model, 10.3% against 2.9% for shuffled communities(34).

**Supplementary Table 3.** The 17 tapped state types, and the structure stream. Symbols are those of Fig. 1. Widths are those of ESMC-300: model width *d* = 960, feed-forward width *f* = 2,560, head width 64. Head-split state types are also probed one head at a time. The structure stream is computed from the stream states rather than tapped inside ESMC. Most state types are a normalization and at most one linear map from the stream. The rotated queries and keys, the attention context and the feed-forward activations are the ones that transform it most.

| State type | Symbol | Family | Description | Width | Heads |
|---|---|---|---|---|---|
| Resid<sub>embed</sub> | *X*<sub>0</sub> | stream | token embeddings, the stream before block 0 | *d* |  |
| Attn<sub>LN</sub> | LN(*X*<sub>*k*</sub>) | attention | attention input after the block's first layer norm | *d* |  |
| Attn<sub>Q</sub>, Attn<sub>K</sub> | *Q*<sub>*h*</sub>, *K*<sub>*h*</sub> | attention | queries and keys after their layer norms, before rotation | *d* | yes |
| Attn<sub>Qrot</sub>, Attn<sub>Krot</sub> | $\widetilde{Q}_h$, $\widetilde{K}_h$ | attention | queries and keys after the rotary embedding | *d* | yes |
| Attn<sub>V</sub> | *V*<sub>*h*</sub> | attention | values | *d* | yes |
| Attn<sub>ctx</sub> | *C* | attention | attention-weighted values, heads concatenated | *d* | yes |
| Attn<sub>out</sub> | *W*<sub>*O*</sub>*C* | attention | output projection, before scaling and the residual add | *d* |  |
| Resid<sub>mid</sub> | *X*<sub>mid</sub> | mid-block | the residual sum between attention and feed-forward network | *d* |  |
| MLP<sub>LN</sub> | LN(*X*<sub>mid</sub>) | feed-forward | feed-forward input after its layer norm | *d* |  |
| MLP<sub>gate</sub>, MLP<sub>up</sub> | *G*, *U* | feed-forward | the two halves of the SwiGLU input projection | *f* |  |
| MLP<sub>act</sub> | *Z* | feed-forward | SiLU(gate) times up, the input of the down projection | *f* |  |
| MLP<sub>out</sub> | *W*<sub>*D*</sub>*Z* | feed-forward | down projection, before scaling and the residual add | *d* |  |
| Resid<sub>post</sub> | *X*<sub>*k*+1</sub> | stream | stream after the block | *d* |  |
| Resid<sub>final</sub> | *X*<sub>final</sub> | stream | output of the final layer norm | *d* |  |
| Struct | *S* | structure | ESMFold2's learned mixture of the stream states, projected to 256 columns | 256 |  |

Two things bound that coverage. Removing every neuron of a tested span moves the prediction past the threshold for only 22.8% of predictions, so for three predictions in four no tested span of blocks is necessary at that token at all. Each prediction is also tested against two candidate communities and not against every community that shares a neuron with it, and with eight candidates, on the first 20,544 held-out predictions, a discovered community explains 4.20% against 0.039% for shuffled ones. A second dictionary of communities, built from each block’s five neurons of largest activation with no gradient, explains 1.65% of predictions against 0.005%, and each of the 193 explaining communities shares neurons with one of its communities, at a median Jaccard index of 0.58.

The 1,022 explained predictions belong to 193 communities with a median of 30 neurons over 9 blocks. Thirty of them account for 80% of the explained predictions, and 27 of those 30 predict one residue for at least 95% of the predictions that they explain. The intact model’s top residue is the true one for 90.0% of explained predictions, against 41.9% of all predictions, and 9.6% of explained predictions are a protein’s first residue, against 0.4% of all, as the initiator methionine has two communities of its own. In each of the 30 most used communities, the most influential neuron removed alone moves the prediction by at most 2.6% of what the whole community does. None of these communities is devoted to a curated term beyond what its residue carries, since when the predictions that a community explains are compared with every other tested prediction of its most predicted residue, none of 51,412 pairs of a community and a curated term is enriched at a false discovery rate of 5% (one-sided Fisher’s exact test, smallest *q* = 0.80), and at most 15% of a community’s predictions carry any one term. With 8 to 124 predictions per community, this test would find only a term that a sizable share of them carries, so it does not exclude a weak preference.

For the concept groups of **Fig. 4G**, 406 of the 413 concepts with a group were tested. A group’s fall exceeds that of its placebo groups by 0.262 nats [0.217, 0.308], and that of each of its ten placebo groups for 81.5% of concepts, while the best single pole exceeds its placebos by 0.0024 nats [0.0001, 0.0049]. The median excess is 0.31 nats for domains, 0.49 for motifs, 0.18 for regions, 0.13 for zinc fingers, and about zero for active sites. The groups are not held out from their own choice, since test F1 chose the blocks and test proteins read the removal, and the placebo groups were chosen on the same proteins.

Two further readings follow Tigges and colleagues(33). When every residue keeps only its largest activations in each block and loses the rest, the model loses far less in its neurons’ basis than in a rotation of it once 64 or more are kept. With 512 of 3,072 kept in all 36 blocks, the KL divergence from the intact model is 0.137 nats and masked-token accuracy falls from 0.397 to 0.355, against 0.77 nats and 0.171 in a rotated basis. The kept neurons are chosen again at every residue, so this cut is not a smaller model. Rebuilding the decoder directions of the SAE on the feed-forward write from the neurons’ write vectors, 13.5% are one neuron’s vector or a sum of up to 64, against 0.14% for rotated neurons, and 86.5% are within no 64, so most of that SAE’s directions are not small sums of neurons.

### Neuron poles: the tests fixed in advance, and readings added later

The main text judges units at residues (**Fig. 5**) because the tests that we fixed in advance over whole proteins could not separate them, and those tests were decided together after the run (**Supplementary Table 4**). Every basis tracks its best term near the AUROC ceiling at both blocks, so the rules on margin and held-out AUROC compare units near it, and the margin, the gap between two near-ceiling scores, is about a thousandth for every basis. At matched firing rates, the median unit’s best term at ESMC-6B’s block 59 reaches a protein-level AUROC of 0.9989 for native poles, 0.9968 to 0.9969 for the ten rotations, 0.9986 for the stream’s dimensions, and 0.9997 for the SAE’s latents, and ESMC-600’s block 26 repeats the pattern. Average precision on the same held-out proteins does not saturate, and it places the neurons above every rotation (0.55 against at most 0.53 in ESMC-600 and 0.59 against at most 0.38 in ESMC-6B) and far below the SAE (0.95 and 0.96) (**Fig. 5D**). The bases differ more in how many units can be scored on held-out clusters, 395 of ESMC-6B’s 13,824 native poles against 67 to 100 for each rotation, 25 of 5,120 stream dimensions, and 1,977 of the SAE’s 16,384 latents. Inheriting the annotations of close homologs, which adds 8,358 positive labels to the 9,324 test proteins, moves held-out AUROC in the fourth decimal and average precision by at most 0.031. Across Biohub’s size and sparsity sweep at ESMC-600’s block 26, the median scored latent’s held-out AUROC stays between 0.998 and 0.999 from 8,192 to 131,072 latents, except with 131,072 latents at *k* = 16 (0.846)(43). A randomly initialized ESMC-6B’s poles track their best terms far less well (median best-term AUROC 0.79) yet with wider margins, so the trained-minus-random margin is negative for five of the seven kinds tested. On the 13 protein-level concept tasks, *k*-sparse probes favor the native axes only where a unit’s extremes are read, since on the products’ maxima and minima at three quarters of the depth, probes on the *k* best native units score above every rotation’s at every *k* from 1 to 16 in both models, while on mean-pooled products, which mix a unit’s two poles, the rotations do at least as well(26). Every rate match at the two blocks was accepted, with Kolmogorov-Smirnov distances of 0.015 to 0.036 (**Methods, “Neuron poles”**), whereas in ESMC-600’s blocks 13 to 16 the median neuron’s gate is open on under 0.02% of residues, so those blocks’ poles cannot reach the SAE’s rates and carry no protein-level comparison.

**Fig. 5B** draws the two SAEs that can be compared across models (**Methods, “Neuron poles”**), and the other sizes that we read keep the order of the units. The comparisons that were fixed in advance read the stream SAE nearest in size to the neurons’ poles, with 8,192 latents for ESMC-600’s 6,144 poles and 16,384 for ESMC-6B’s 13,824, and an SAE of 65,536 latents at ESMC-6B’s block 59. The stream SAE nearest in size to ESMC-600’s 6,144 poles, with 8,192 latents, identifies 238 concepts at block 26, against 178 for the neurons and 290 for the 16,384-latent SAE, and an SAE of 65,536 latents identifies 368 at ESMC-6B’s block 59, against 262 for the 16,384-latent one, so larger dictionaries identify more concepts and every SAE that we read identifies more than the neurons. ESMC-6B, read at every fourth block, repeats ESMC-600’s profile over depth (**Supplementary Fig. 5A**). By feature type, the neurons and the SAE on the feed-forward write share most domains and regions, and the SAE alone reads most of the motifs, zinc fingers, and active sites that either reads (**Supplementary Fig. 5D**).

A paired test, added after the comparison and not among the rules fixed in advance, reads each concept’s best unit, chosen on validation, by its test F1. Under the neurons that F1 exceeds the one under a rotation by a median of 0.07 to 0.08 in ESMC-300 (3 rotations), 0.11 to 0.13 in ESMC-600 (10 rotations), and 0.16 to 0.18 in ESMC-6B (10 rotations), over the 631 to 636 concepts scored under both, and a two-sided Wilcoxon signed-rank test rejects equality for each of the 23 rotations at a Holm-corrected *p* below 10*^−^*^55^(73, 75).

Under InterPLM’s code, ESM2-8M’s SAEs reach 2,512 associated latents in a layer and its residual dimensions 51, close to the published 2,548 and 46, and InterPLM’s permuted control identifies two concepts or fewer at every layer(22). In ESM2-8M’s first three layers the dimensions have the larger share of associated units, though they identify fewer concepts (at layer 3, 51 of 320 dimensions on 9 concepts against 216 poles on 74), so the comparison fixed in advance holds at 9 of the 12 layers.

Cammarata judged the neurons of vision models clean by a further sign, that a search for the rotation under which the fewest units carry each input’s activity came back to the neurons in every block(32). In ESMC the same search, run from three random starts, ends nearly as far from the neurons as it began. At ESMC-600’s block 26 it finds rotations 11 to 20% sparser than the neurons by its own measure, yet these identify 89 to 102 concepts, more than the rotations they started from (58 to 62) but about half as many as the neurons (**Fig. 5B**), so sparsity alone does not explain the neurons’ lead. At ESMC-6B’s block 59 the search stalls far short of the neurons’ sparsity, and its rotations identify 48 to 54 concepts, about as many as their starts.

Three readings were added after the residue-level comparison, following the report of Tigges and colleagues that, in a language model, a neuron’s meaning narrows as it fires harder where a random direction’s does not, that a neuron often reads along the write of one earlier neuron, and that transcoder features copying no single neuron are mostly groups of 2 to 64 neurons that fire together(33). Each was read against rotations by a rule written before it ran, and none takes a decision. For firing strength, each unit’s residues at three quarters of each model’s depth were cut by its own activation into four bands, from the top 5 to 1% of residues to the top 0.01%, and a band’s precision is the share of its test residues inside the concept that covers most of its validation residues. The neurons’ median rise in precision from the weakest band to the strongest exceeds that of each of three rotations at ESMC-600’s block 26 (0.018 against 0.011 to 0.012) but not at ESMC-300’s block 22 (0.023 against 0.023 to 0.025) or at ESMC-6B’s block 59 (*−*0.004 against 0.043 to 0.044), and only 1.2, 1.1, and 1.0% of the poles have a strongest band mostly inside one concept, so narrowing with strength is confined in ESMC to a small tail of poles.

For wiring, read from the weights alone, a neuron’s link is the largest absolute cosine between its gate or up weights, scaled by the layer norm’s gain, and the write vector of any neuron of the block before, and a link is strong above 0.3, as in that report(36). At all 143 pairs of adjacent blocks in the three models, more neurons have a strong link to a neuron’s write than to an axis of a rotation of those writes, with 11.1, 8.2, and 17.1% of gate weights and 6.0, 4.3, and 2.4% of up weights strongly linked in ESMC-300, ESMC-600, and ESMC-6B, although at eight pairs of ESMC-6B 44 to 73% of gates also link to rotated writes, so there the 0.3 mark does not single out neurons. For groups of poles, each latent of the 131,072-latent SAEs on the feed-forward write that fires in both halves of the dictionary test’s 2 million residues was fit with 1 to 64 poles of its block, chosen greedily on one half, and the fit’s correlation with the latent was read on the other (**Supplementary Fig. 5B**). Fits of 64 rotated poles track latents that carry 3.6 and 1.5% of the SAE’s activity at ESMC-600’s block 26 and ESMC-6B’s block 59, against 33.0 and 38.7% for the neurons’ poles, and for the concepts that only the SAE reads the 64-pole fits reach a median held-out correlation of 0.10 and 0.12. A single pole already tracks latents that carry a quarter to a third of that SAE’s activity, and many transcoder features of language models are likewise single neurons(95). These fits use activity alone, whereas Tigges and colleagues counted groups from a transcoder’s decoder weights, so the two counts are not the same measure.

**Supplementary Table 4.**
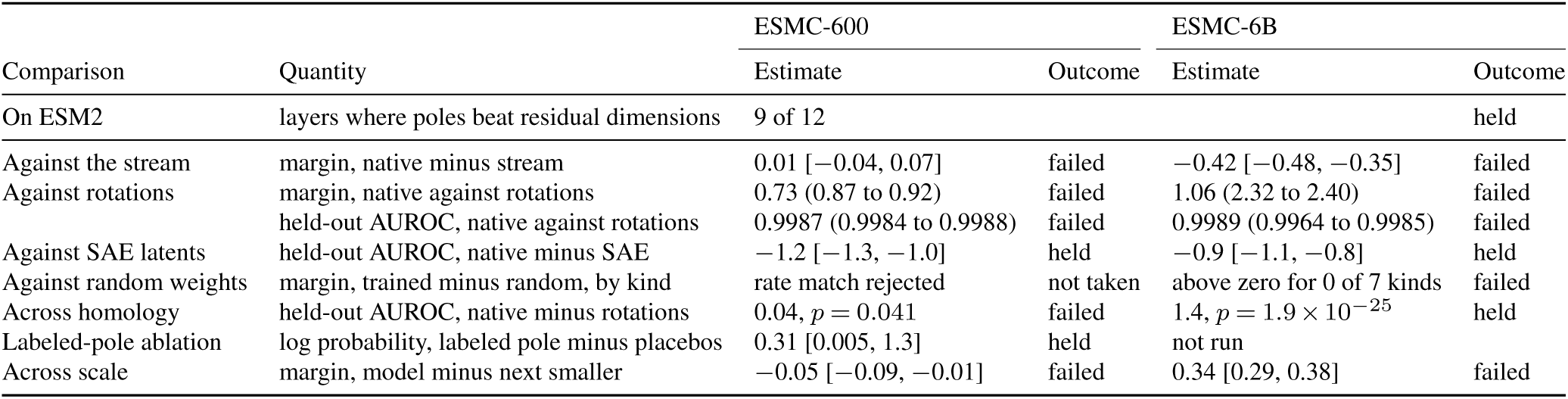
The decisions fixed in advance. For each comparison fixed before the run, the quantity its rule reads, the estimate in each model and the outcome, at ESMC-600’s block 26 and ESMC-6B’s block 59. Margins, AUROC differences and the fall in log probability after a labeled pole is ablated (nats) are in thousandths (*×*10*^−^*^3^), with 95% intervals over units in brackets (90% for the comparison with SAE latents). Against the rotations, the native poles’ value is followed in parentheses by the range of the ten rotations, and the AUROC row is not rescaled. Margins and held-out AUROC are medians over units at the SAE’s firing rates. The comparison with random weights covers every block of ESMC-600, where the rate match was rejected at every block but the first, the *p* values across homology are one-sided, the ablation covers 85 poles, and for scale each column is that model minus the next smaller one. The comparison on InterPLM’s pipeline was decided on ESM2-8M and ESM2-650M(5). Reading the neurons’ products by sign alone takes the same decision at every row, with estimates within 0.0001, and so does deciding the comparisons with SAE latents and across homology again after the test proteins inherit their close homologs’ labels.

| Comparison | Quantity | ESMC-600 |  | ESMC-6B |  |
|---|---|---|---|---|---|
|  |  | Estimate | Outcome | Estimate | Outcome |
| On ESM2 | layers where poles beat residual dimensions | 9 of 12 |  |  | held |
| Against the stream | margin, native minus stream | 0.01 [−0.04, 0.07] | failed | −0.42 [−0.48, −0.35] | failed |
| Against rotations | margin, native against rotations | 0.73 (0.87 to 0.92) | failed | 1.06 (2.32 to 2.40) | failed |
|  | held-out AUROC, native against rotations | 0.9987 (0.9984 to 0.9988) | failed | 0.9989 (0.9964 to 0.9985) | failed |
| Against SAE latents | held-out AUROC, native minus SAE | −1.2 [−1.3, −1.0] | held | −0.9 [−1.1, −0.8] | held |
| Against random weights | margin, trained minus random, by kind | rate match rejected | not taken | above zero for 0 of 7 kinds | failed |
| Across homology | held-out AUROC, native minus rotations | 0.04, *p* = 0.041 | failed | 1.4, *p* = 1.9 × 10<sup>−25</sup> | held |
| Labeled-pole ablation | log probability, labeled pole minus placebos | 0.31 [0.005, 1.3] | held | not run |  |
| Across scale | margin, model minus next smaller | −0.05 [−0.09, −0.01] | failed | 0.34 [0.29, 0.38] | failed |

**Supplementary Fig. 5.**
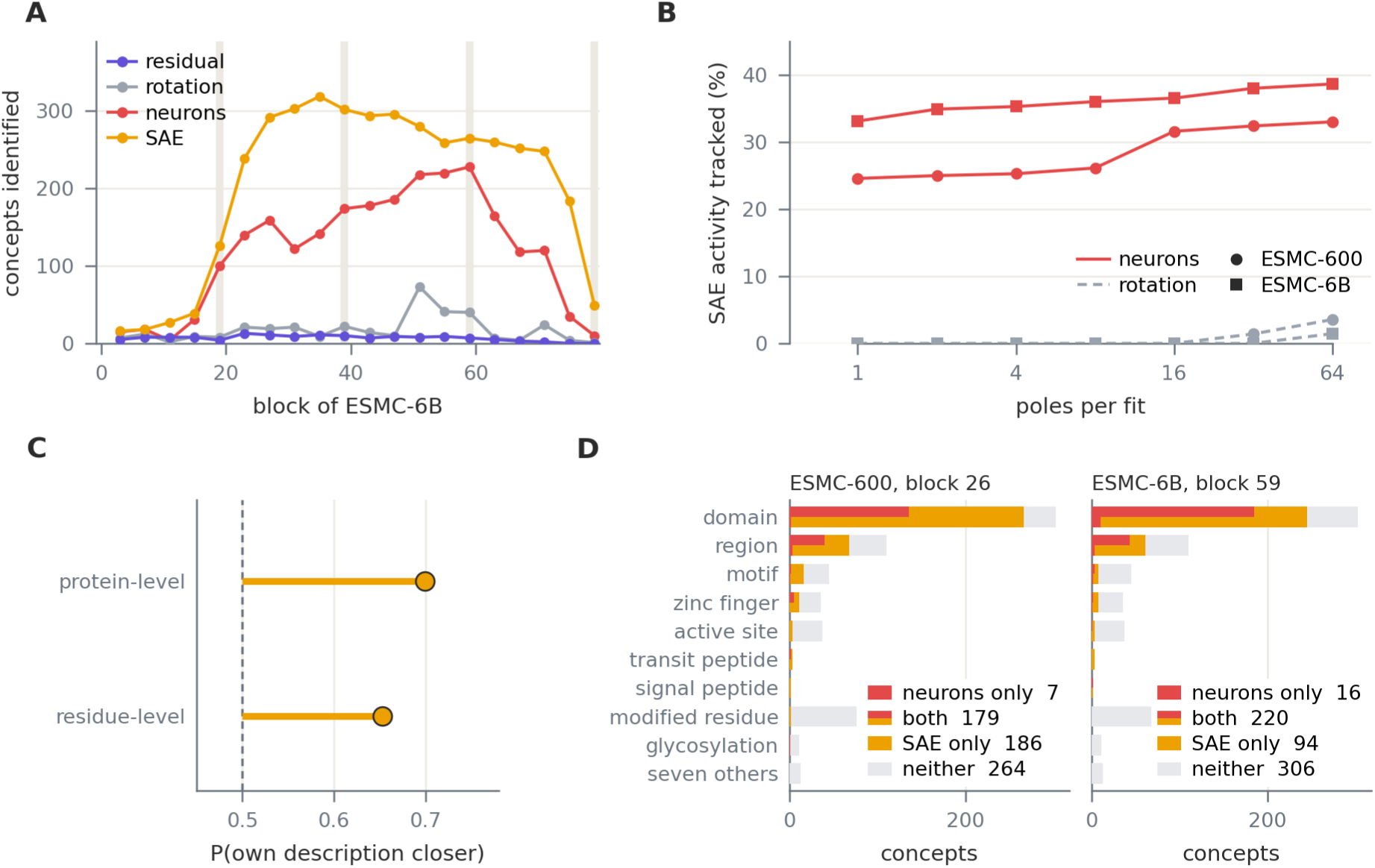
Further readings of the units. **A**, concepts, of 636, identified at every fourth block of ESMC-6B by the stream’s dimensions, one random rotation, the neurons, and each block’s SAE on the stream (16,384 latents), with the four blocks where the comparison was planned shaded. **B**, the share of the activity of Biohub’s SAE on the feed-forward write (131,072 latents) that lies on latents which a fit of the neurons’ poles tracks on held-out residues, and the same for a rotation’s poles, as the fit may use more poles. A single pole tracks a quarter to a third of the activity, groups of up to 64 poles add little, and the fits track the latent for 1 of the 186 and 2 of the 94 concepts that only the SAE reads. **C**, for the latents of Biohub’s described SAE of ESMC-6B, the probability that a latent’s curated protein-level or residue-level label is closer, by word overlap, to its own multi-agent description than to a description matched for category and length (0.70 and 0.65, against 0.5 by chance, dashed line). Both exceed every one of 1,000 permutations. **D**, of 636 Swiss-Prot concepts, those of each feature type that the neurons’ best pole reads, that the best latent of the SAE on the feed-forward write reads, that both read, or that neither reads, at three quarters of each model’s depth. A concept counts here when its unit, chosen on validation proteins, has an F1 above 0.5 on test proteins.

### A curated label needs a testable term, and is not evidence of use

The main text names a unit by the curated term it tracks best (**Fig. 5E**), and what such a label means is bounded by the dictionary on which the labeling was first run and by the poles whose use we tested. Before the neuron study, we labeled every latent of Biohub’s described ESMC-6B SAE (layer 60, 16,384 latents, *k* = 64) with a structured annotation vocabulary and compared the labels with the descriptions that Biohub generated for the same latents(23, 24, 43, 44). The atlas covers 575,503 Swiss-Prot proteins (release 2026_02) and 73,980 terms, 7,013 of them local, each carried by at least ten proteins(35). A latent’s global score for a protein is its maximum activation, tested against each protein-level term by tie-corrected Mann-Whitney AUROC and by positive-activation enrichment, and its local score is the within-protein difference between the shares of active residues inside and outside a term’s intervals, with a latent-term pair selected at nominal Benjamini-Hochberg *q ≤* 0.01 in one family over every testable endpoint of every latent(76). Every latent received qualifying global and local labels, 9,568 endpoints per latent on average, and randomized controls show how little that coverage says. With annotation rows shuffled across proteins, 496 to 509 of a panel of 512 latents still received a label in each of 20 replicates, shuffling only within strata of taxonomic domain by length octave labeled every latent and reproduced 42% of the observed endpoints, and circular shifts of each protein’s activation track labeled every latent with 13% of the observed local endpoints. Whether a latent has a label therefore separates nothing, and taxonomy and length alone reproduce two fifths of the global breadth, which is why the main text counts a label only when it is confirmed on held-out proteins.

The labels agree with Biohub’s descriptions above matched chance (**Supplementary Fig. 5C**), as judged by one global and one local representative per latent, frozen before any comparison and scored against the latent’s own description and against descriptions permuted within strata of Biohub category by word count. In that first labeling, by word TF-IDF similarity, the own description scored above a matched one with a mean win probability of 0.7165 for global representatives and 0.5975 for local ones, above all 1,000 permutations (*p* = 1*/*1001), and two embedding similarities agreed(96–100). In a blinded audit of 192 latents, language-model raters judged 58.1% of own pairs biologically corresponding against 29.7% of matched pairs, and 171 of the 223 correspondences shared related context rather than a compatible concept, so the two procedures point at the same neighborhood of biology more often than at the same concept.

The labels predict their terms weakly across sequence clusters when every unannotated test protein counts as a negative, as representatives reselected on training proteins reached a median test AUROC of 0.5307 over 11,967 global representatives, on test clusters that had been examined during development. Relabeled on the neuron study’s compendium and atlas set, where a latent is scored on test proteins only when its best term has ten positive and ten negative test proteins observed in its aspect, 1,977 latents qualify, with a median held-out AUROC of 0.9999 and average precision of 0.963 against a median prevalence of 0.00013, and agreement with Biohub’s descriptions holds at win probabilities of 0.699 and 0.653 (both *p* = 1*/*1001; **Supplementary Fig. 5C**). Whether a label generalizes therefore depends first on whether its term can be tested at all.

To test whether the model uses what a labeled pole reads, we took the 85 of 200 locally labeled poles of ESMC-600’s block 26 that had enough held-out carriers (**Supplementary Fig. 6A**). Ablating each where it fires inside its term, by setting its product to the unit’s mean where it does not fire, lowers the masked tokens’ log probability by a median of 0.00031 nats [0.000005, 0.0013] more than ablating its ten placebo poles of matched activation mass, and leaves masked tokens outside the term no worse. The test removes one pole at a time, and the poles that share a concept’s label cost far more when they are removed together (**Supplementary Information, page 24**). Removing each unit’s weakest 75% of activations in every block raises the masked-token loss on 68,573 masked tokens of 1,000 held-out proteins from 1.935 to 2.081 nats and lowers accuracy from 0.397 to 0.353, while removing its strongest activations of the same total magnitude, 5.9% of them, raises the loss to 3.199 nats and cuts accuracy to 0.095 (**Supplementary Fig. 6B**). Steering a masked token’s stream along the curve that its hydropathy context, the mean Kyte-Doolittle value of its flanking residues(101), traces at block 26 moved its prediction no more naturally than steering along a straight line (path energy 0.3632 against 0.3637), in a weak test whose largest offsets moved the predicted hydropathy by about 0.1(102) (**Supplementary Fig. 6C**). A linear probe on ESMC-600’s pooled stream after block 26 splits exactly into one term per neuron of blocks 0 to 26 and the rest(103). Over 754 Gene Ontology and InterPro targets those neurons carry a median 40% of the probe’s separation, while the neurons labeled with a target carry a median share of 0.000 and at most 0.122, above matched unlabeled neurons for 62 of 218 targets(37, 63) (**Supplementary Fig. 6D**).

### Tuning and the choice of state

The main text reads interaction at a state chosen on validation with a tuned probe (**Fig. 6C**), and on the 2024 gold standard the tuning and the state add separately. We fitted a tuned linear probe and a tuned MLP to each state and to the one-hot encoding (**Methods, “Protein-protein interaction”**). At ESMC-6B’s MLP_up_ of block 77, our survey’s probe reproduces its archived score (test MCC 0.293 against 0.293), and tuning lifts it by 0.017 *±* 0.003 ROC AUC with an MLP and 0.009 *±* 0.002 with a linear probe. The state inside the block leads the stream state under every probe, by 0.037 *±* 0.009, 0.049 *±* 0.009, and 0.035 *±* 0.009 ROC AUC under our survey, tuned linear, and tuned MLP probes, and the one-hot encoding gains nothing from tuning. The last hidden state behaves the same way in all three models, reaching under tuned MLPs an MCC of 0.239, 0.249, and 0.244 and a test ROC AUC of 0.672 *±* 0.016, 0.675 *±* 0.015, and 0.674 *±* 0.016 for ESMC-300, ESMC-600, and ESMC-6B. The best stream states of the two smaller models already add 0.024 *±* 0.009 and 0.019 *±* 0.011 over it under tuned MLPs, and the state inside the block (MLP_gate_ of block 28 and of block 32) only 0.006 *±* 0.008 and 0.011 *±* 0.008 beyond them, against 0.035 *±* 0.009 in ESMC-6B, so reading inside the block adds clearly only in the largest model. **Fig. 6C** draws our rows without their intervals, which are *±*0.014 to 0.016 in ROC AUC and *±*0.016 to 0.018 in PR AUC for every row except random vectors (*±*0.007 and *±*0.011), and the tuned probes’ MCC was not bootstrapped.

A one-hot encoding reads this dataset well above chance (**Fig. 6C**), so part of what pooled-state probes read here is not interaction itself, in line with an audit that found substantial partner-independent signal in protein-disjoint benchmarks through SAE fingerprints and with our earlier finding that sampling choices let pLM-based predictors learn localization or taxonomy in place of interaction(10, 66, 104).

**Supplementary Fig. 6.**
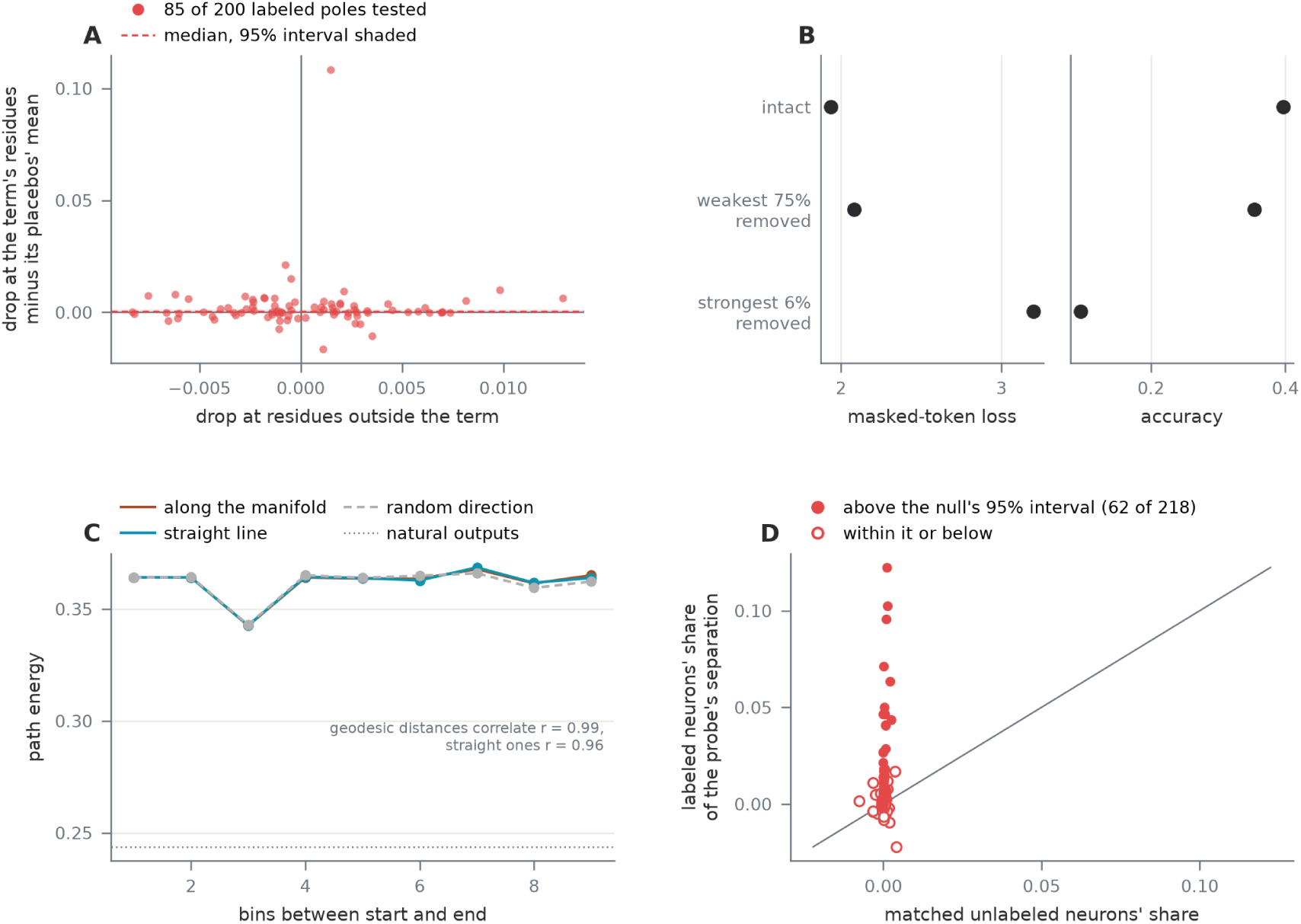
How much ESMC-600 uses what labeled poles read. **A**, for each of the 85 labeled poles of block 26 with enough held-out carriers, the fall in the masked tokens’ log probability when the pole is set to its non-firing mean wherever it fires inside its term, minus the mean fall for its ten placebo poles of matched activation mass, against its own fall at masked tokens outside the term. The dashed line is the median, its 95% interval shaded. **B**, masked-token loss and accuracy over 68,573 masked tokens of 1,000 held-out proteins with the model intact, with each unit’s weakest 75% of activations removed in every block, and with its strongest activations of the same total magnitude removed (6% of activations). **C**, median path energy, the Bhattacharyya distance from the predicted distribution to the behavior manifold averaged over a path’s steps, by the number of hydropathy bins between start and end, for steering along the fitted manifold, along the straight line between the same centroids and along a random direction of the same length; the dotted line is the unsteered outputs’ energy. **D**, for each of the 218 probe targets with a labeled neuron, the labeled neurons’ share of the probe’s separation on test proteins against the mean share of matched unlabeled neurons. Filled points lie above the matched sets’ 95% interval, and the line marks equality.

### Settings, controls, and datasets

The Methods point to the three tables of this section, which list the global training settings, the controls with their targets, and the datasets with their sources and sizes, and the paragraphs below give the settings that the Methods summarize and the compute that the experiments took.

The 18 UniProt datasets are split by MMseqs2 clusters. Clusters at 20% identity (30% for active site, binding site, conserved site, and repeat) go whole to training plus validation or to test, and validation clusters are drawn at 50% identity from the rest, with 80% coverage of the target sequence(90). The four BioMap tasks keep their benchmark test splits, in which NetSurfP-2.0’s secondary structure test chains share at most 25% identity with training and localization is partitioned at 30% identity(105, 106). Their validation sets are a random 10% of training, so they carry no homology guarantee between training and validation. The interaction gold standard shares no protein between its splits, whose sequences were made nonredundant with CD-HIT at 40% identity(45, 107, 108).

The sweep behind **Supplementary Table 5** trained every probe of five state types under 14 settings, learning rates from 10*^−^*^4^ to 10*^−^*^1^ in half-decade steps by a weight decay of 0 or 0.1, at two batch sizes for each regime. A probe’s shortfall under a setting is its best validation score over all settings minus its score under that setting, and among the settings within 0.005 of the lowest mean shortfall the one whose sweep finished fastest was taken. ProtParam was trained under its own setting, since under the classification setting its probes fell 0.29 short of their best validation scores on average. Protein probes train for at most 100 epochs and stop after 10 validations without a lower validation loss, and residue probes for at most 10 epochs and stop after 3.

The converged refits fit the same loss on the same standardized training rows. Classification probes use L-BFGS along 14 L2 penalties from 10*^−^*^6^ to 3.2, each run until its largest absolute gradient fell below 10*^−^*^4^, ProtParam uses ridge regression in closed form, and all 26 fits of **Supplementary Table 2** converged(68). MLP probes were swept over the linear probes’ learning rates and weight decays, hidden widths of 256, 512, and 1,024, and dropout of 0.1 or 0.3. Classification MLPs took a hidden width of 256, dropout of 0.1, a learning rate of 10*^−^*^3^, weight decay of 0.1, and batches of 256 proteins, and ProtParam’s MLPs a hidden width of 512, dropout of 0.3, a learning rate of 3 *×* 10*^−^*^4^, weight decay of 0.1, and batches of 1,024.

All experiments ran in containers, the survey, the MLP study, the probe variants, the geometry, and the interventions on NVIDIA GH200 GPUs with 96 GB of memory, and the familiarity analysis and the tuned pair probes on NVIDIA H100 GPUs. Writing one model’s pooled stores took a few minutes for ESMC-300 and about a quarter of an hour for ESMC-6B, and training and scoring every probe of one residue task under all controls took up to about an hour. The MLP study, with its sweeps, three seeds, and bootstrap intervals, took about a day of GPU time, the tuned pair probes a few GPU-hours, and the familiarity analysis about one. The neuron study ran on a GH200, where reading its 83,190 proteins once took 51 minutes through ESMC-600 and 1 hour 57 minutes through ESMC-6B, and the tests of communities, concept groups, sparsity, and erasure levels in ESMC-600 took six and a half hours on another.

**Supplementary Table 5.** Global training settings. , chosen on validation scores of the sweeps. Mean shortfall is the average gap between a probe’s validation score under the setting and under its own best setting, so one setting per regime costs about 0.03 at most.

| Probes | Batch | Learning rate | Weight decay | Epochs | Patience | Mean shortfall |
|---|---|---|---|---|---|---|
| Classification (12 tasks) | 256 proteins | 3 × 10<sup>−2</sup> | 0 | 100 | 10 | 0.027 |
| Regression (ProtParam) | 1,024 proteins | 10<sup>−3</sup> | 0.1 | 100 | 10 | 0.002 |
| Residue (9 tasks, 5 swept) | 32,768 tokens | 3 × 10<sup>−3</sup> | 0.1 | 10 | 3 | 0.033 |

**Supplementary Table 6.** Controls and their targets. Each control removes one source of signal (sequence order, learned weights, a trained readout, state norms, per-protein variation or the input itself), so a probe’s lead over a control measures what that source contributes.

| Control | Input | Targets |
|---|---|---|
| Scrambled sequence | every residue after the first shuffled, seeded per sequence | ProtParam recomputed on the scrambled sequence, residue labels and multilabel vectors shuffled within the protein by an independent draw, a single class kept |
| Untrained model | states of a copy of ESMC-300 reset to PyTorch defaults(109) | unchanged |
| Untrained probe | the probe’s initial weights | unchanged |
| Unit-norm tokens | every token state scaled to unit L2 norm before pooling or probing | unchanged |
| Batch mean | the mean state of each batch of 8 test proteins (position by position for residue tasks) | the batch’s mean (regression) or most common label |
| Gaussian input | standard normal states | unchanged |

**Supplementary Table 7.** Datasets. Source names the database, and citations give the database paper and the paper the dataset came from. Labels counts classes (multiclass), labels (multilabel) or targets (regression). Split gives the fractions of proteins in training, validation and test (for pair tasks, of the proteins in each split’s pairs). Homology split states what keeps test proteins apart from training proteins, and the pair dataset shares no protein between splits. The InterPro binding and active site tasks, with under 750 proteins each, are where the choice of probe matters most.

| Dataset | Level | Type | Labels | Proteins | Split | Homology split | Source |
|---|---|---|---|---|---|---|---|
| ProtParam | protein | regression | 31 | 18,126 | 0.70/0.15/0.14 | MMseqs2, 20% | UniProt(14, 35, 62) |
| InterPro binding site | protein | multilabel | 39 | 640 | 0.70/0.14/0.17 | MMseqs2, 30% | UniProt, InterPro(14, 35, 37, 110) |
| InterPro active site | protein | multilabel | 58 | 718 | 0.67/0.16/0.17 | MMseqs2, 30% | UniProt, InterPro(14, 35, 37, 110) |
| InterPro repeat | protein | multilabel | 65 | 1,189 | 0.69/0.16/0.15 | MMseqs2, 30% | UniProt, InterPro(14, 35, 37, 110) |
| InterPro conserved site | protein | multilabel | 334 | 2,948 | 0.70/0.15/0.15 | MMseqs2, 30% | UniProt, InterPro(14, 35, 37, 110) |
| InterPro family | protein | multilabel | 239 | 3,601 | 0.58/0.19/0.23 | MMseqs2, 20% | UniProt, InterPro(14, 35, 37, 110) |
| InterPro domain | protein | multilabel | 364 | 8,003 | 0.66/0.17/0.17 | MMseqs2, 20% | UniProt, InterPro(14, 35, 37, 110) |
| InterPro homologous superfamily | protein | multilabel | 100 | 8,198 | 0.67/0.17/0.16 | MMseqs2, 20% | UniProt, InterPro(14, 35, 37, 110) |
| GO molecular function | protein | multilabel | 123 | 11,504 | 0.70/0.16/0.15 | MMseqs2, 20% | UniProt(14, 35, 63) |
| GO biological process | protein | multilabel | 304 | 12,255 | 0.70/0.15/0.14 | MMseqs2, 20% | UniProt(14, 35, 63) |
| GO cellular component | protein | multilabel | 164 | 16,443 | 0.69/0.15/0.16 | MMseqs2, 20% | UniProt(14, 35, 63) |
| Metal ion binding | protein | binary | 1 | 7,332 | 0.74/0.08/0.18 | not reported | BioMap (PDB)(14, 61, 111, 112) |
| Localization | protein | multiclass | 10 | 7,530 | 0.71/0.08/0.22 | DeepLoc, 30% | BioMap (UniProt)(14, 61, 106) |
| Lipidation | residue | multiclass | 25 | 978 | 0.67/0.13/0.20 | MMseqs2, 20% | UniProt(14, 35) |
| Topology | residue | multiclass | 17 | 4,777 | 0.71/0.16/0.14 | MMseqs2, 20% | UniProt(14, 35) |
| Peptide | residue | multiclass | 4 | 4,977 | 0.70/0.14/0.16 | MMseqs2, 20% | UniProt(14, 35) |
| Functional sites | residue | multiclass | 6 | 6,893 | 0.69/0.16/0.15 | MMseqs2, 20% | UniProt(14, 35) |
| Phosphorylation | residue | multiclass | 6 | 8,648 | 0.70/0.15/0.15 | MMseqs2, 20% | UniProt(14, 35) |
| Secondary structure | residue | multiclass | 4 | 8,986 | 0.70/0.15/0.15 | MMseqs2, 20% | UniProt(14, 35) |
| SSP3 | residue | multiclass | 3 | 11,479 | 0.85/0.09/0.06 | NetSurfP-2.0, 25% | BioMap (PDB)(14, 61, 105, 112) |
| SSP8 | residue | multiclass | 8 | 11,479 | 0.85/0.09/0.06 | NetSurfP-2.0, 25% | BioMap (PDB)(14, 61, 105, 112) |
| Post-translational modification | residue | multiclass | 12 | 15,349 | 0.70/0.16/0.15 | MMseqs2, 20% | UniProt(14, 35) |
| PPI, Bernett, et al. 2024 | pair | binary | 1 | 11,019 | 0.39/0.34/0.27 | CD-HIT, 40% | HIPPIE v2.3(45, 113) |

